# Denoised MDS-UPDRS Part-III Scores Yield New Patterns of Progression Heterogeneity in Early Stage Parkinson’s Disease

**DOI:** 10.64898/2026.05.04.722810

**Authors:** Jonathan D. Koss, Sule Tinaz, Hemant D. Tagare

## Abstract

Parkinson’s Disease (PD) Motor Scores (MDS-UPDRS Part III) are quite noisy. This paper proposes a new methodology for processing these scores by first denoising the scores to enhance the underlying progression signal, and then conducting a high-dimensional analysis which does not sum the scores into a total movement score. The analysis gives novel insights into PD progression heterogeneity: it reveals that the heterogeneity is continuously variable rather than clustered into “subtypes” and that the variability is along two easily understood axes. This analysis also resolves some of the discrepancies in previously reported progression subtypes. Finally, the analysis reveals that patient-specific progression cannot be predicted from baseline using only MDS-UPDRS Part III scores.

## 1 Introduction

The progression of Parkinson’s disease (PD) is known to be heterogeneous, i.e., the trajectory of the disease varies significantly from patient to patient. A quantitative understanding of this heterogeneity is important because it can shed light on the etiology of PD, predict PD progression at the individual level, and inform cohort selection and outcome monitoring in PD clinical trials.

PD heterogeneity is usually analyzed using PD subtypes. Each subtype represents a specific pattern of disease progression. Many different approaches for defining subtypes are used in the literature. One approach, based on clinical phenomenology, defines the following subtypes: tremor dominant (T-D), postural instability and gait disorder dominant (PIGD-D) [1–3], and akinetic-rigid dominant (AR-D) [4–6]. A shortcoming of the subtyping approach is that subtypes do not seem to be stable. PD patients appear to change subtypes throughout the course of the disease, even over the course of a few years [7–10]. One explanation offered for this change is that subtypes represent different stages of PD progression rather than differences in the underlying disease trajectories [7, 11–13]. Other subtyping approaches stratify PD by features such as age at onset [14–18], genetics [19–24], or the predominant non-motor symptom [25].

Data-driven subtypes obtained through cluster analysis have also been proposed [8, 26–28]. However, clusters identified by such methods are not consistent with each other, with some methods finding clusters that match T-D vs PIGD-D/AR-D subtypes while others finding clusters consistent with late-onset, fast progression and early-onset, slow progression [29]. Yet other clustering methods suggest novel subtypes such as mild-motor predominant and diffuse malignant [30–32].

From a data-science point of view, the methods summarized above have some limitations: First, in many cases, subtypes are defined cross-sectionally rather than longitudinally, e.g. [29, 33–35]. That is, they are defined by the state of the disease at a fixed point in time instead of examining the longitudinal trajectory of the disease.

Second, most subtypes are defined using motor scores, which tend to be very noisy. This noise often masks the underlying progression of the disease, and can cause correlations between various observations to be smaller. Noise can also cause spurious clustering.

Third, motor scores are also ordinal rather than numerical, and some statistical care is needed to deal with ordinality. Previous clustering-based attempts to produce subtypes do not account for ordinality of the scores nor do they use noise models that are appropriate for ordinal data [27, 31, 32, 35].

Fourth, many (but not all) subtypes are defined by summing motor scores for different symptoms into a single Total Movement Score (TMS) [3, 9, 26, 36]. Using TMS obscures differential progression in the canonical motor symptoms (tremor, bradykinesia, rigidity, etc.). This is problematic because differential progression is likely to be critical to subtyping.

Finally, subtyping effectively places a patient in one of a finite number of categories without allowing for continuous gradation between categories [8]. But whether a finite category model of progression heterogeneity is more realistic than a continuously variable heterogeneity model is a priori unclear.

The goal of this study is to develop a new methodology for analyzing PD heterogeneity in motor scores using longitudinal scores while taking the above critique into account. We are especially interested in denoising the motor scores before drawing statistical conclusions. Denoising the scores improves their signal-to-noise ratio, and thus improves the reliability of statistical analysis. We denoise the Movement Disorder Society’s Unified PD Rating Scale (MDS-UPDRS) Part III scores using a recently reported method which accounts for the ordinal nature of the scores [37]. We also do not sum motor scores into a TMS. Instead, we analyze the multivariate time series of all MDS-UPDRS Part III scores.

Visualizing the multivariate time series of MDS-UPDRS Part III scores (and the accompanying statistical analysis) shows the absence of clear clusters in the data. Given this, we propose a methodology to understand how the progression data are continuously distributed. The methodology also finds and uses neighborhoods within which the data can be statistically summarized. A new understanding of PD progression heterogeneity emerges from this analysis.

In addition to quantifying heterogeneity, a goal of this paper is to examine whether longitudinal scores can be predicted from baseline. In data-science terms, a time series which is predictable from a fixed point in time is *Markovian*, else it is *non-Markovian*. Thus we are interested in assessing whether the denoised time series of MDS-UPDRS Part III scores is Markovian or non-Markovian. The data for our analysis come from the Parkinson’s Progression Markers Initiative (PPMI), a longitudinal study with the goal of finding biomarkers for PD progression (www.ppmi-info.org) [38].

All PPMI subjects provided a signed informed consent form. The data was gathered in accordance with Institutional Review Board/Independenct Ethics Commitee approval. All data is available in anonymized form.

All PPMI patients were recruited at initial PD diagnosis. We refer to this time point as the baseline. Details about inclusion and exclusion criteria for PPMI can be found in [38]. MDS-UPDRS scores are collected by PPMI for each patient at 3-6 month intervals over the first 5 years and then annually. However, missing visits are common. MDS-UPDRS scores are collected in an off-medication state.

We selected patients from PPMI who had at least 5 visits in the time span from baseline to baseline plus four years. This resulted in 386 patients being selected. Of these 96% were white, 2% were Asian, and the remaining participants were black, native American, or Pacific Islanders. They had a mean of 11.9 (± 3.3) visits. Their mean age was 61.2 (±9.7) years and 62% were male. At baseline, the mean Hoehn and Yahr (H&Y) score was 1.8 (±0.4) and the mean MDS-UPDRS Part III total movement score (TMS) was 21.3 (±7.7).

Of the 386 patients, 66 (17.1%) had pathogenic *LRRK2* variants and 20 (7.5%) had pathogenic *GBA* variants, while 84 (21.8%) had *APOE ε4* variants. DaTscan images were available at baseline and baseline plus year four for 198 patients.

## 2 Results

Because we chose not to analyze motor scores by summing them into a TMS, and because we do not wish to make any prior assumptions about the existence of well-formed clusters in the data, our analysis pipeline is quite different from analysis pipelines reported in the literature. Our pipeline is illustrated in Figure 1. Figure 1a illustrates our ideas geometrically, while Figure 1b illustrates the processing and analysis as a flow chart. Each step of Figure 1b is summarized below.

**Figure 1:**
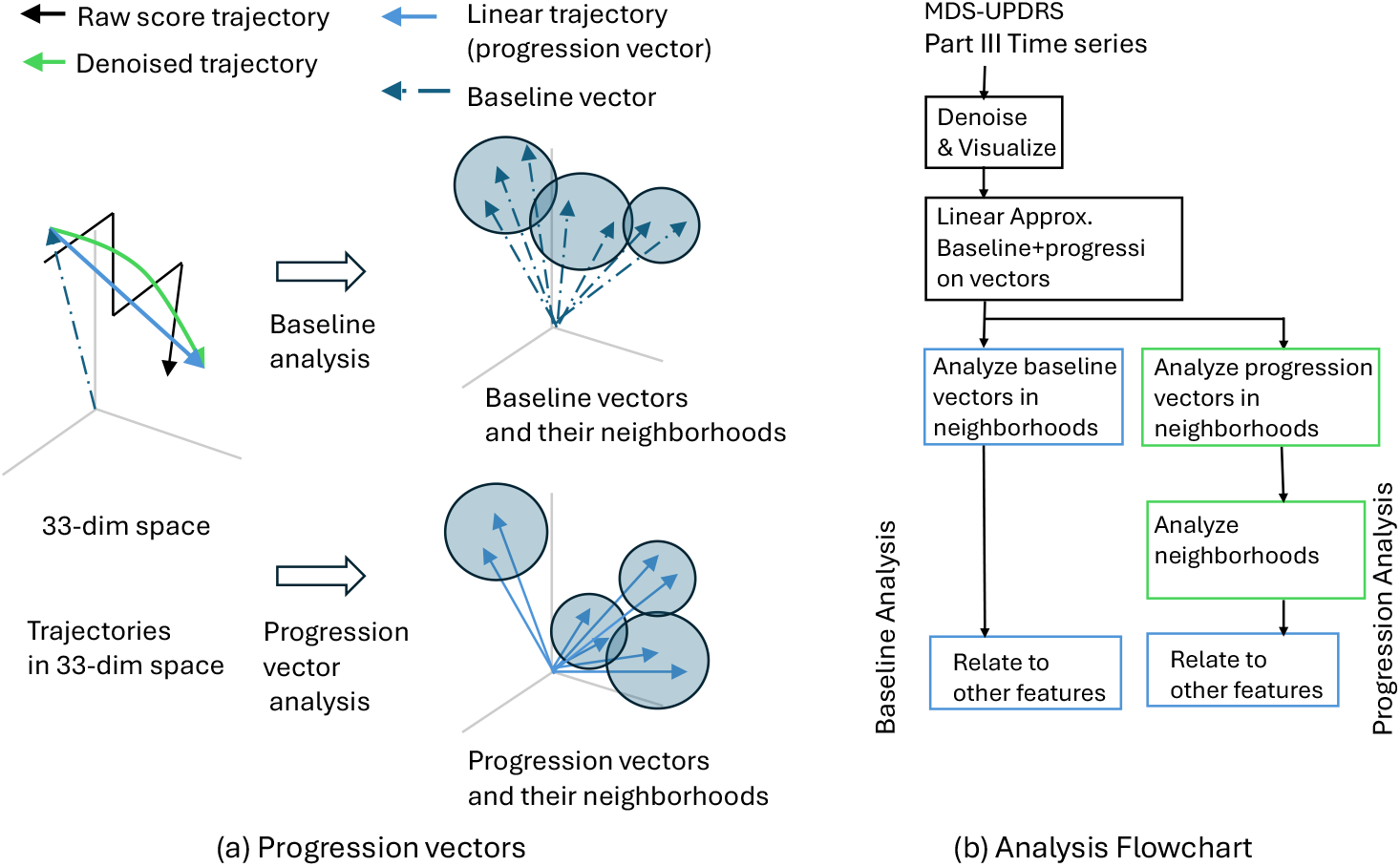
Overview of the analysis. (a) Longitudinal 33-dimension MDS-UPDRS (Movement Disorder Society-Sponsored Unified Parkinson’s Disease Rating Scale) scores for a PD (Parkinson’s disease) patient form a noisy curve in a 33-D (33-dimension) space (solid-black trajectory). Denoised scores give a smooth trajectory (solid-green trajectory), which in turn can be efficiently approximated by a linear trajectory (linear-blue trajectory) from baseline to baseline plus year four. The trajectory is analyzed in terms of its baseline vector (dashed arrow) and progression vector (blue arrow). Baseline analysis is carried by statistical analysis of neighborhoods in the vectors are distributed. Progression analysis is carried by moving all progression vectors so that their tail is at the origin, and then statistically analyzing neighborhoods in which the vectors are distributed.(b) Flowchart of the analysis steps.

We begin by considering all MDS-UPDRS Part III scores, except the Hoehn-Yahr score, for every participant. The Hoehn-Yahr score is omitted because it represents an overall summary of the PD state, rather than a specific movement feature. The other 33 scores are movement specific. For each patient, at any point in time, the 33 scores are arranged in a 33-dimensional (33-D) vector. As the disease progresses, this vector sweeps out a (noisy) curve in the 33-D space of scores (Figure 1a, solid black trajectory). Each such curve represents the progression of the disease in a single patient.

We denoise the 33 scores to obtain a 33-D denoised progression curve for each patient (Figure 1a, solid green trajectory). We refer to the pre-denoised scores as the *raw scores* and the post-denoised scores as the *denoised scores*. Thus, we obtain a single raw score trajectory and a single denoised score trajectory per patient.

Next, we visualize the 33-D trajectories by using multidimensional scaling (MDS) to embed the 33-D data into a 2-D space. Each 33-D data point from every trajectory is embedded as a point in the 2-D space. Connecting the embedded points from each trajectory gives an embedding of the trajectory in the 2-D space.

As discussed in detail in Section 2.2 below, the above MDS visualization reveals that the raw trajectories are extremely noisy and significantly mask the underlying progression. Progression is more clearly revealed by the denoised trajectories. Denoised trajectories have two properties: First, the denoised trajectories are almost linear, and second, the trajectories are non-Markovian (they cannot be predicted from the baseline). Consequently, we separately analyze each trajectory by decomposing it into its baseline vector (Figure 1a, dashed vector from origin) and the changes from the baseline vector to the end (Figure 1a, solid blue vector). We call the latter the *progression vector*. To be precise, suppose *y*_*i*_, *i* = 1, *…, T* is the trajectory in the 33-D space, then the baseline vector is *y*_1_, and the progression vector is *y*_*T*_ − *y*_1_. The progression vector informs about the total progression of the disease from baseline.

After decomposing each trajectory into a baseline vector and progression vector we visualize the spread (distribution) of these vectors in 33-D space. Visualization clearly shows that the vectors do not cluster in any obvious way (Section 2.2). Additional statistical analysis of the baseline and projection vectors shows that their distributions are not multimodal. That is, denoised baseline and projection data appear to be spread continuously without any evidence of a natural clustering. Because there is no natural clustering, we adopt the following strategy to to statistically summarize the continuous spread of baseline and progression vectors: We find compact spherical neighborhoods in the 33-D space such that the neighborhoods cover all data (Figure 1a). The neighborhoods are found by hierarchically clustering the vectors and finding a level in the hierarchy where the spheres seem to compactly fit the data. Methodological details of this step are in Section 4.9. We emphasize that we do not take the clusters to represent subtypes. We simply consider each cluster to be a local data neighborhood within which the data is reasonably homogeneous so that local summarizing statistics can be calculated. To emphasize that the clusters are not subtypes but simply local regions for statistical analysis, we refer to them as *data neighborhoods*, or more simply as *neighborhoods*.

Baseline and progression vectors are both subjected to neighborhood analysis. Progression vectors are analyzed by translating them so that their tail is at the origin.

The vectors in each neighborhood are summarized statistically by grouping the scores into Bradykinesia and Rigidity subscores (B+R) (sum of MDS-UPDRS Part III scores: 3.2, 3.3, 3.4, 3.5, 3.6, 3.7, 3.8 (18 scores)), Tremor subscores (sum of MDS-UPDRS Part III scores 3.14, 3.15, 3.16, 3.17, 3.18 (10 scores), and Axial subscores (sum of MDS-UPDRS Part III scores 3.9, 3.10, 3.11, 3.12 (4 scores)). Bar plots of the means of these canonical subscores provide a summary of the data in each neighborhood. Statistical comparison of these subscores in different clusters provides an understanding of heterogeneity.

We further analyze the spread of the progression vectors by investigating the locations of their spherical neighborhoods in terms of the distance and the direction of the centers of the neighborhood.

Finally, we relate the statistics of baseline and progression vectors to other patient features such as demographics, genetic mutations, and DaTscan images. Each of the above steps is discussed in detail in Figure 1.

### 2.1 Denoising

As mentioned above, we begin with the raw MDS-UPDRS Part III score time series for each patient (Figure 1a) and denoise the time series of each score using the method reported in [37] and described in detail in the Methods Section 4.3 of this paper. This denoising method takes into account the ordinality of data and the short duration of the time series. The method is intrinsically capable of handling irregularly spaced time series, so data interpolation for missing visits is not required. The method uses a hierarchical model to draw statistical power from the entire cohort of PPMI patients while denoising each patient-specific time series.

We use this method to denoise MDS-UPDRS Part III scores from baseline to baseline plus 4 years (examples shown in Figure 2). During denoising, scores for bi-lateral questions are re-arranged by the more/less-affected side instead of left/right (described in Methods Section 4.4).

**Figure 2:**
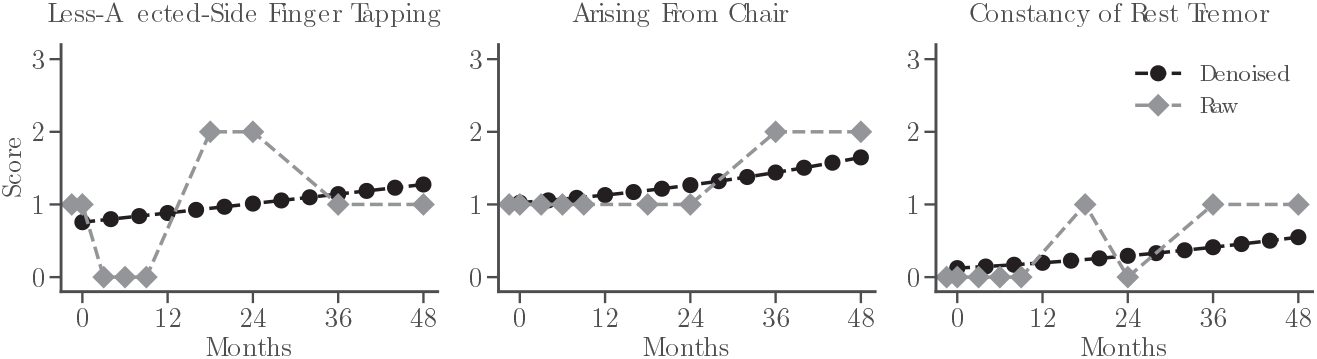
Examples of raw and denoised time series of some MDS-UPDRS (Movement Disorder Society-Sponsored Unified Parkinson’s Disease Rating Scale) Part III (three) scores.

We note here that the publication [37] extensively validated the denoising method with simulated data that mimicked MDS-UPDRS Part III scores and with actual MDS-UPDRS Part III scores. The method was also compared with traditional smoothing. The publication [37] shows that the method outperforms traditional techniques, works well with simulated MDS-UPDRS part III data, and does not overfit actual MDS-UPDRS part III data.

Denoising can affect the signal along with the noise. To assess how much the method affects the underlying signal, we performed an additional principal component analysis (PCA) of the raw and denoised data (see Supplementary Information). The first few principal components of noisy signals are usually sensitive to the signal, and a comparison of these shows that they are very similar for raw and denoised data. This suggests that denoising does not substantially change the underlying signal in the data.

### 2.2 Visualization with Multidimensional Scaling

Figure 3a shows the multidimensional-scaling embedding of the raw (non-denoised) 33-D MDS-UPDRS Part III trajectories to a plane (2-D) for 30 randomly chosen patients. Clearly, the raw trajectories are very noisy. Any attempt to statistically analyze these trajectories is likely to be strongly influenced by the noise. Figure 3b shows the denoised trajectories of the same patients. Disease progression is now more clearly evident.

**Figure 3:**
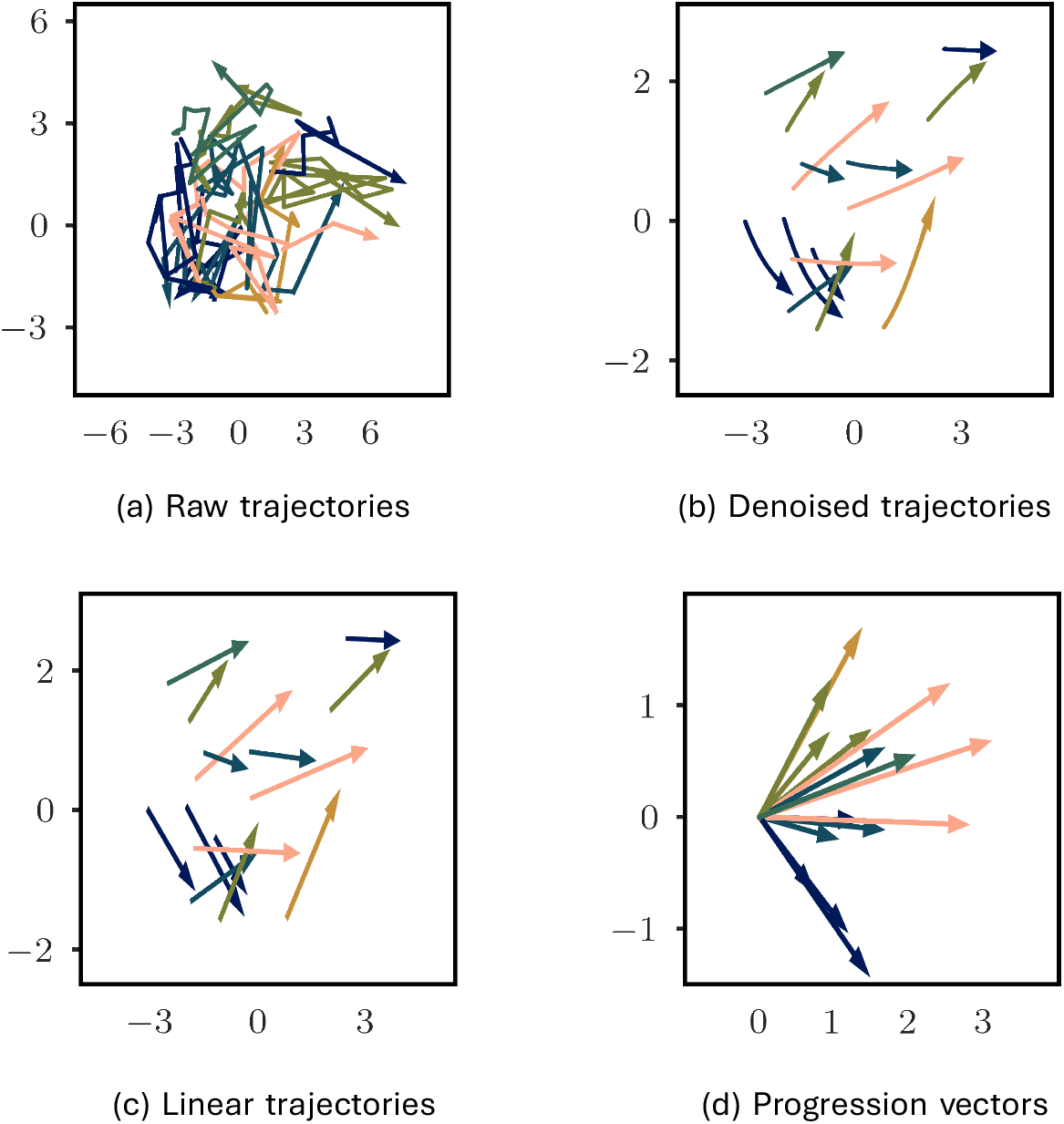
Visualizing sample trajectories by embedding them in 2-dimensions with multidimensional scaling. (a) 15 sample raw trajectories. (b) Denoised trajectories. (c) Linear approximation of the denoised trajectories. (d) Progression vectors of the linear approximation.

Figure 3b also suggests that the denoised trajectories can be reasonably approximated as linear. Hence, we took the first and last time points of the denoised trajectories to create *linear trajectories* shown in Figure 3c. Linear trajectories can be analyzed in terms of their baselines and progression vectors. The baseline vector is the vector from the origin to the denoised baseline score. The progression vector is the vector from the denoised baseline score to the baseline plus year 4 score. It represents the change in the scores over this span of time (examples shown in Figure 3d). Because we analyze data over a fixed time period of 4 years, the progression vector can also be thought of as a rate of progression over this span of time.

### 2.3 Analysis of Denoised Baseline Scores

Figure 4a shows the 33-D baseline scores of all subjects projected to 2-D using multidimensional scaling. Each point (dot) in the figure is the projection of a baseline score of a single patient. The colored contours indicate density of points obtained via a kernel-density estimate (described in Section 4.6). The contours of the kernel-density estimate suggest that baseline scores are unimodal. Unimodality implies that the heterogeneity at baseline is continuously variable and does not exhibit distinct clusters. Well-defined subtypes would imply a multimodal density.

**Figure 4:**
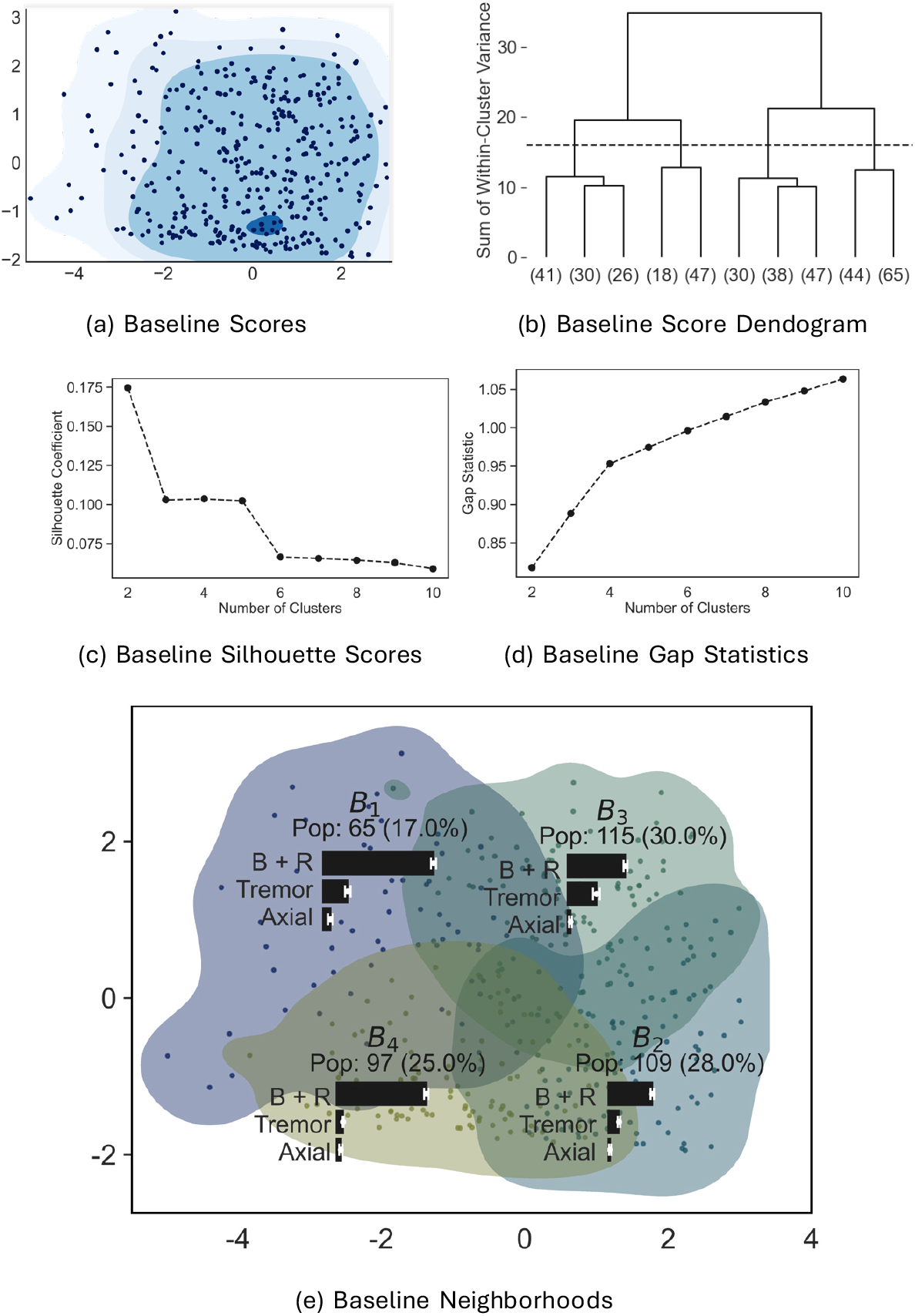
Visualization and hierarchical clustering of denoised baseline scores in 33-dimension space of MDS-UPDRS (Movement Disorder Society-Sponsored Unified Parkinson’s Disease Rating Scale) Part III (three) scores. (a) Scatter plot of baseline scores with density projected onto 2-dimensions using multi-dimensional scaling. (b) Dendrogram: four clusters (indicated by the dashed line) were chosen as data neighborhoods (referred to as *B*_1_, …, *B*_4_); (c) Silhouette Scores; (d) Gap Statistics. (e) Baseline neighborhoods projected on to 2-D. The population in each neighborhood is indicated. The mean (±stdev.) motor subscores of each neighborhood are shown as bar graphs. B+R (bradykinesia plus rigidity); Pop. (population)

### 2.4 Data Modality

Motivated by Figure 4a we statistically evaluated baseline data for unimodality. The classic statistical test of unimodality is Hartigan’s dip-test [39]. However, this test only applies to univariate data. In the literature, several have attempted to extend Hartigan’s dip-test to multivariate data, but none is generally accepted. In one approach, called the *dip-dist* test [40], the scalar Euclidean distances between each data point and all other data points are calculated and tested for unimodality using Hartigan’s dip-test (Methods Section 4.7 for a longer description). Accepting the null hypothesis of unimodality of distances at each point amounts to testing for unimodality of the data. In another approach, called the *folding test* [41], multivariate data are “folded” back on themselves and the original variance of the data is compared with the folded variance to test for unimodality (see Methods Section 4.7).

We applied the dip-dist test and the folding test to the baseline scores. The dip-dist test, with Bonferroni and Benjamini-Yekutieli corrections for multiple testing, applied to the 33-D denoised baseline scores of 386 patients could not reject the null hypothesis of unimodality at *p* = 0.05. The folding test also accepted the hypothesis of unimodality at *p* = 0.0001 (Φ = 5.10). We also carried out additional tests of unimodality vs multi-modality the SigClust simulation method (available in R). The result, which is available in the Supplemenatry Material, also accepts the hypothesis of unimodality.

Finally, we evaluated whether there were any linearly separable clusters in the data. This was done by projecting the 33-D data repeatedly onto 1-D with 10, 000 random Gaussian projections. Rejecting the null hypothesis of uni-modality in any projection would indicate the presence of linearly separable clusters in the data. We applied the Hartigan dip-test with Bonferroni and Benjamini-Hochberg corrections for multiple testing to evaluate unimodality of the projection. We were unable to reject unimodality in any of the projections at *p* = 0.05.

### 2.5 Neighborhoods for analyzing data

After establishing that the 33-D denoised baseline scores were unimodal, we sought to characterize the heterogeneity by creating different neighborhoods within to summarize the population. To generate these neighborhoods, we hierarchically clustered the baseline scores using Agglomerative Clustering (see Section 4.9 for details). To select a number of clusters, we looked at a dendrogram drawn with Ward linkage method, silhouette scores, and the gap statistics for different numbers of clusters (see Figure 4b-d). These three statistics did not agree on a clear choice of the number of clusters which is expected as the data was shown to be unimodal (see Section 2.4). We chose four clusters as a compromise between having sufficiently many neighborhoods to capture heterogeneity, while keeping the neighborhoods large enough to give reliable statistics, and being a reasonable choice of clusters according to all three metrics (see Section 4.9).

We label these neighborhoods *B*_1_, …, *B*_4_. The multidimensional scaling projections of the neighborhoods along with the projection of the data points is shown in Figure 4c. The overlap in the *B*_1_,…, *B*_4_ neighborhoods in the 2-D visualization occurs because the clustering is performed on the full 33-D baseline scores, but projected in 2-D. Figure 4c also shows the population of *B*_1_, *…, B*_4_. The neighborhoods *B*_2_, *B*_3_ contain the majority of the data (58%).

### 2.6 Heterogeneity in Baseline Motor Scores

Figure 4c shows the mean baseline B+R, Tremor and Axial subscores in each neighborhood as bar graphs. These graphs suggest that the subscore means are different in each neighborhood. To assess this more formally, we calculated the means and standard deviations of TMS, H&Y, and B+R, Tremor, and Axial subscores of baseline data in *B*_1_, …, *B*_4_ (displayed as the BL rows in Table 1). We used two-sided t-tests (unequal variances, unequal data sizes) to assess whether the means were similar. After adjusting for multiple comparisons (Benjamini-Hochberg correction), none of the hypotheses of equality of means were significant at *p* = 0.05. This suggests that each of *B*_1_, *…, B*_4_ contain baseline scores with distinct mean values, confirming heterogeneity at baseline.

**Table 1:** Comparison of mean TMS, B+R, Tremor, Axial, and H&Y scores in baseline neighborhoods *B*_1_, …, *B*_4_. Each of TMS, B+R, Tremor, Axial parts, and H&Y have two rows. The BL row shows the mean and std. dev. of the score for the neighborhood. The Δ row shows the mean and std. dev. of the score progression vector for the neighborhood. Bold-faced neighborhoods below each entry indicate the neighborhood that has a significantly different mean.

| | | $B_1$ | $B_2$ | $B_3$ | $B_4$ |
| --- | --- | --- | --- | --- | --- |
| B+R | BL | 23.03(0.53) | 9.58(0.34) | 12.41(0.4) | 18.82(0.45) |
|  |  | **B1 B2 B3 B4** | **B1 B2 B3 B4** | **B1 B2 B3 B4** | **B1 B2 B3 B4** |
| | $\Delta$ | 4.96(2.69) | 5.13(2.05) | 5.4(2.77) | 5.02(2.76) |
|  |  | **B1 B2 B3 B4** | **B1 B2 B3 B4** | **B1 B2 B3 B4** | **B1 B2 B3 B4** |
| Tremor | BL | 5.74(0.56) | 2.88(0.26) | 6.55(0.69) | 2.02(0.14) |
|  |  | **B1 B2 B3 B4** | **B1 B2 B3 B4** | **B1 B2 B3 B4** | **B1 B2 B3 B4** |
| | $\Delta$ | 0.78(1.39) | 0.89(1.3) | 1.17(1.34) | 0.77(1.16) |
|  |  | **B1 B2 B3 B4** | **B1 B2 B3 B4** | **B1 B2 B3 B4** | **B1 B2 B3 B4** |
| Axial | BL | 2.24(0.46) | 1.07(0.21) | 1.31(0.27) | 1.59(0.33) |
|  |  | **B1 B2 B3 B4** | **B1 B2 B3 B4** | **B1 B2 B3 B4** | **B1 B2 B3 B4** |
| | $\Delta$ | 1.07(0.84) | 0.82(0.67) | 0.89(0.76) | 0.95(0.86) |
|  |  | **B1 B2 B3 B4** | **B1 B2 B3 B4** | **B1 B2 B3 B4** | **B1 B2 B3 B4** |
| TMS | BL | 31.8(6.05) | 13.9(4.07) | 20.75(4.87) | 23.12(4.78) |
|  |  | **B1 B2 B3 B4** | **B1 B2 B3 B4** | **B1 B2 B3 B4** | **B1 B2 B3 B4** |
| | $\Delta$ | 7.0(3.91) | 7.05(2.91) | 7.63(3.89) | 6.92(3.71) |
|  |  | **B1 B2 B3 B4** | **B1 B2 B3 B4** | **B1 B2 B3 B4** | **B1 B2 B3 B4** |
| H&Y | BL | 1.88(0.32) | 1.43(0.38) | 1.54(0.36) | 1.8(0.31) |
|  |  | **B1 B2 B3 B4** | **B1 B2 B3 B4** | **B1 B2 B3 B4** | **B1 B2 B3 B4** |
| | $\Delta$ | 0.23(0.29) | 0.35(0.28) | 0.37(0.24) | 0.23(0.23) |
|  |  | **B1 B2 B3 B4** | **B1 B2 B3 B4** | **B1 B2 B3 B4** | **B1 B2 B3 B4** |

### 2.7 Progression vectors in *B*_1_, *…, B*_4_

Next we analyzed how baseline scores relate to progression scores. To this end, we compared the mean progression vectors for TMS, B+R, Tremor, and Axial subscores in the neighborhoods *B*_1_,…, *B*_4_ (displayed as the Δ rows in Table 1) with a two-sample t-test. None of the null hypotheses of equality of means could be rejected at *p* = 0.05 after adjusting for multiple comparisons (Benjamini-Hochberg correction). This suggests that progression in all baseline neighborhoods was similar over 4 years. That is, knowing the baseline neighborhood does not contribute to predicting the progression vector.

### 2.8 Predicting Progression Vectors

We further explored whether baseline vectors can predict progression vectors by using Partial Least Squares Regression (PLS) and Principal Component Regression (PCR). The goal of these analyses is to evaluate whether the entirety of the progression vector, i.e. each of its 33 components, can be estimated from the entirety of the baseline vector.

PLS was used to estimate each component of the progression vector from the entire 33-D baseline vector. When analyzing a progression vector component, the baseline and progression vector (component) data was split into a training (80%) and cross-validation (20%) set. The PLS model with *k* components was used to fit the training data set, and the learned model was used to predict the progression vector component using the cross-validation set. The *R*^2^ (coefficient of determination) value of the test set prediction was used to asses the quality of the prediction. The *R*^2^ value is the fraction of the progression vector component variance explained by PLS. Values of *R*^2^ less than 0.2 are generally taken to indicate no or poor prediction.

The number of PLS components (*k*opt) which gave the highest *R*^2^ value for the cross-validation set was chosen, and are reported in Table 2. Next, the entire was dataset was repeatedly repartitioned into a training (80%) and test set (20%) for 1000 times. For each partition, PLS was fit to the training set using *k*opt components, and evaluated on the test set using *R*^2^. Table 2 shows the median ± median-absolute-deviation values of *R*^2^ for the training and test set over the 1000 trials for each progression vector component. As Table 2 shows, the largest *R*^2^ value for the test set occurs for the fourth component (Arm rigidity on the less affected side). This value, and in fact all *R*^2^ values for the test data, are less than 0.2, many significantly so, suggesting no or very poor ability to predict the progression vector from the baseline.

**Table 2:** Partial Least Squares Regression (PLS) and Principal Component Regression (PCR) of denoised baseline vector with each component of denoised progression vector. The first column shows the identity of the progression vector component. Within PLS and PCR, the first column shows the optimal number of components obtained by cross-validation, and the second and third columns show the R^2^ value of the regression for training and test data. The largest R^2^value for Test data occurs for the 4th progression vector component for PLS and the 26th component for PCR. Both are displayed in bold font.

|  | PLS |  |  | PCR |  |  |
| --- | --- | --- | --- | --- | --- | --- |
| | Num.<br>Comp.<br>($k_{\text{opt}}$) | $R^2$ Train<br>med (mad) | $R^2$ Test<br>med (mad) | Num.<br>Comp.<br>($k_{\text{opt}}$) | $R^2$ Train<br>med (mad) | $R^2$ Test<br>med (mad) |
| 1 | 3 | 0.145(0.058) | -0.014(0.058) | 1 | 0.003(0.013) | -0.012(0.013) |
| 2 | 4 | 0.156(0.055) | -0.013(0.055) | 1 | 0.007(0.015) | -0.004(0.015) |
| 3 | 2 | 0.166(0.056) | 0.020(0.056) | 4 | 0.001(0.012) | -0.014(0.012) |
| 4 | 3 | 0.291(0.055) | **0.156(0.055)** | 3 | 0.015(0.019) | -0.003(0.019) |
| 5 | 1 | 0.176(0.052) | 0.136(0.052) | 4 | 0.106(0.038) | 0.090(0.038) |
| 6 | 3 | 0.137(0.062) | -0.037(0.062) | 1 | 0.010(0.019) | -0.005(0.019) |
| 7 | 2 | 0.172(0.069) | 0.023(0.069) | 4 | 0.017(0.018) | 0.003(0.018) |
| 8 | 1 | 0.153(0.048) | 0.107(0.048) | 3 | 0.032(0.024) | 0.016(0.024) |
| 9 | 2 | 0.150(0.048) | 0.034(0.048) | 2 | 0.061(0.030) | 0.046(0.030) |
| 10 | 1 | 0.098(0.044) | 0.045(0.044) | 1 | 0.026(0.024) | 0.010(0.024) |
| 11 | 1 | 0.058(0.036) | 0.011(0.036) | 4 | 0.024(0.023) | 0.007(0.023) |
| 12 | 1 | 0.068(0.039) | 0.014(0.039) | 1 | 0.010(0.019) | -0.007(0.019) |
| 13 | 4 | 0.248(0.073) | 0.065(0.073) | 4 | 0.066(0.031) | 0.047(0.031) |
| 14 | 1 | 0.060(0.036) | -0.006(0.036) | 2 | 0.006(0.015) | -0.010(0.015) |
| 15 | 4 | 0.240(0.064) | 0.069(0.064) | 3 | 0.039(0.026) | 0.022(0.026) |
| 16 | 1 | 0.042(0.031) | -0.020(0.031) | 1 | 0.008(0.017) | -0.009(0.017) |
| 17 | 4 | 0.160(0.056) | -0.024(0.056) | 4 | 0.007(0.015) | -0.008(0.015) |
| 18 | 1 | 0.145(0.057) | 0.079(0.057) | 3 | 0.070(0.032) | 0.049(0.032) |
| 19 | 4 | 0.264(0.078) | 0.073(0.078) | 2 | 0.002(0.012) | -0.014(0.012) |
| 20 | 2 | 0.248(0.139) | -0.005(0.139) | 4 | 0.020(0.023) | 0.007(0.023) |
| 21 | 2 | 0.309(0.127) | 0.173(0.127) | 4 | 0.037(0.026) | 0.014(0.026) |
| 22 | 1 | 0.037(0.034) | -0.034(0.034) | 2 | 0.006(0.016) | -0.008(0.016) |
| 23 | 4 | 0.305(0.065) | 0.140(0.065) | 4 | 0.024(0.024) | 0.010(0.024) |
| 24 | 3 | 0.136(0.051) | -0.019(0.051) | 4 | 0.044(0.031) | 0.031(0.031) |
| 25 | 3 | 0.215(0.069) | 0.082(0.069) | 4 | 0.044(0.034) | 0.027(0.034) |
| 26 | 4 | 0.261(0.079) | 0.097(0.079) | 4 | 0.008(0.017) | -0.011(0.017) |
| 27 | 2 | 0.225(0.073) | 0.106(0.073) | 4 | 0.073(0.037) | **0.053(0.037)** |
| 28 | 2 | 0.132(0.048) | 0.019(0.048) | 4 | 0.025(0.022) | 0.009(0.022) |
| 29 | 2 | 0.087(0.046) | -0.011(0.046) | 3 | 0.001(0.011) | -0.012(0.011) |
| 30 | 1 | 0.087(0.042) | 0.030(0.042) | 3 | 0.037(0.028) | 0.024(0.028) |
| 31 | 4 | 0.138(0.073) | -0.068(0.073) | 4 | 0.007(0.017) | -0.010(0.017) |
| 32 | 4 | 0.088(0.068) | -0.281(0.068) | 1 | 0.002(0.015) | -0.016(0.015) |
| 33 | 2 | 0.081(0.042) | -0.050(0.042) | 4 | 0.003(0.013) | -0.011(0.013) |

We also analyzed the relation between baseline vectors and progression vectors using PCR. The motivation for using PCR is that the components of baseline vectors are likely to be correlated, and this correlation can adversely affect the regression of baseline and progression vectors. PCR uses principal component analysis to eliminate the correlation and provide a more robust regression. We evaluated PCR in the same manner as we evaluated PLS – an initial training-cross-validation split and the *R*^2^ value was used to find the optimal number of components. Then, as above, a 1000-fold repartitioning of the data was used to calculate the *R*^2^ values of for training and test data for each progression component. The results are also shown in Table 2. The progression vector component with the highest *R*^2^ value is “hand kinetic tremor on the less-affected side”. This *R*^2^ value, and in fact all *R*^2^ values, are significantly less that 0.2, suggesting no or very poor ability to predict the progression vector from the baseline.

### 2.9 Other differences in *B*_1_, *…, B*_4_

In a further attempt to investigate what was different about the baseline neighborhoods, we compared age, sex, genetic mutations and DaTscan striatal binding ratios (SBR, calculated as described in Section 4.12) in *B*_1_, …, *B*_4_. Table 3 shows the mean values (±stdev.) for age, the male/female ratio, ratios of pathogenic mutations for genetics, and mean (±stdev.) baseline SBR (BL row of DaTscan entry) and SBR change in (baseline-year 4, Δ row of DaTscan entry) in each neighborhood. Statistical tests (two-sided t-tests with differing variance and counts for age and SBR baseline and change, two sample Z-tests for equality of proportions for male/female ratio and genetic mutations) with Benjamini-Hochberg correction for multiple comparisons showed that only pathogenic *LRRK2* mutations were significantly different. The occurrence of the mutations in *B*_2_, *B*_4_ had a significantly higher frequency than in *B*_1_, *B*_3_.

**Table 3:**
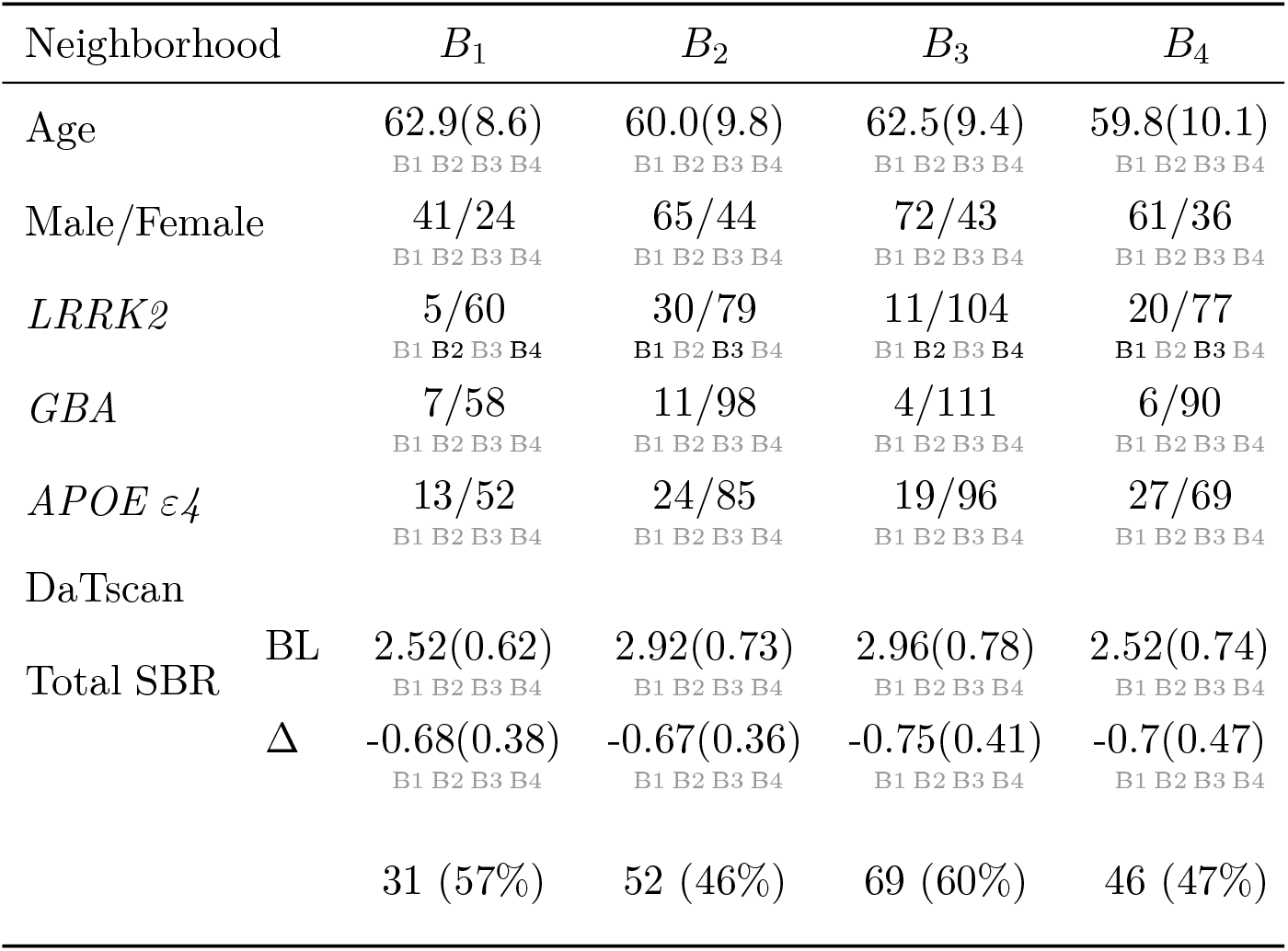
Comparison of age, sex, genetics, and DaTscan striatal binding ratios (SBR)s for baseline neighborhoods. For numerical variables (age, SBR): means and standard deviations in each neighborhood are shown. For genetics, the numbers indicate YES/NO counts. For *LRRK2* and *GBA*, YES indicates the presence of a pathogentic mutation. For *APOE ε4*, YES indicates the presence of the *APOE ε4* allele. Bold-faced neighborhoods below a value indicate that they are significantly different from the above value.

### 2.10 Summary of baseline results

In summary, the following picture emerges for the heterogeneity of denoised baseline scores:

1. Denoised baseline scores do not show evidence of clustering. They seem to be continuously spread in the data space.
2. Denoised baseline scores are heterogeneous. Different regions of the data space, as represented by the different baseline neighborhoods, have statistically different subscores.
3. Denoised baseline scores do not or very poorly predict progression.
4. *LRRK2* mutations have a higher rate in *B*_2_, *B*_4_. However, there are no differences in age, Male/Female ratios, other genetic mutations, and SBRs.

### 2.11 Heterogeneity of Progression Vectors

Next, we analyzed progression vectors. This analysis was carried out by translating all progression vectors so their tails were at the origin. Then multidimensional scaling was used to visualize the tips of the progression vectors, see Figure 5a. Figure 5a is analogous to Figure 4a; each point (dot) in Figure 5a represents the progression vector of a participant. The colored contours represent the density of points (i.e. density of progression vectors). The baseline scores, progression vectors are spread continuously rather than being clustered.

**Figure 5:**
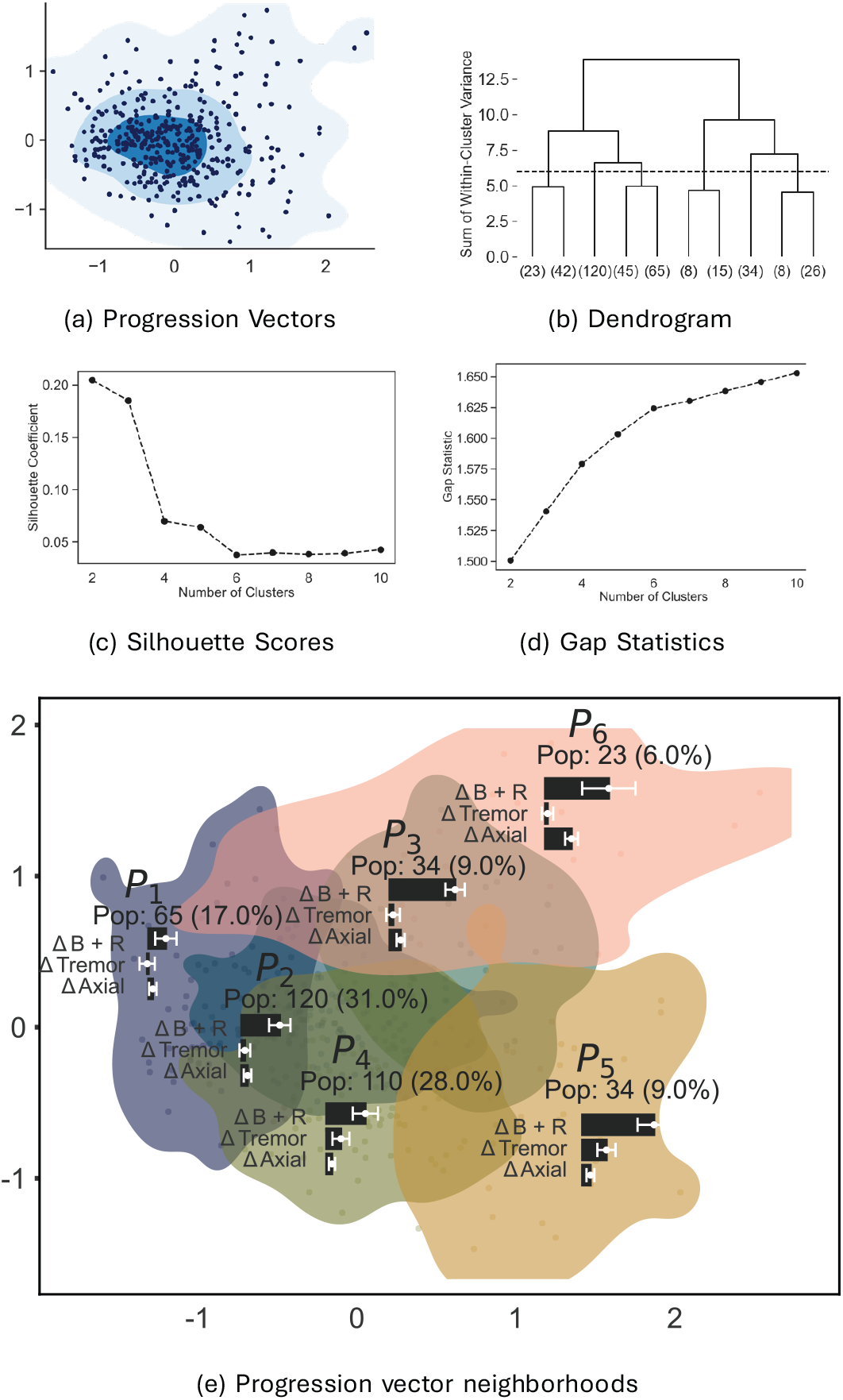
Visualization of denoised progression vectors in 33-D (33-dimensional) space of MDS-UPDRS (Movement Disease Society-Sponsored Unified Parkinson’s Disease Rating Scale) Part III (three) scores (a) Two dimensional scatter plot and density of progression vectors after multidimensional scaling. (b) Dendrogram; (c) Silhouette Scores; (d) Gap Statistics. (e) Six neighborhoods in which progression vectors are visualized in 2-D (2-dimensions) after multidimensional scaling. The bar graphs in each neighborhood show mean progression of B+R (bradykinesia plus rigidity), Tremor, and Axial features. Pop. (population)

### 2.12 Data Modality

We tested for unimodality of progression vectors using the dip-dist and folding tests mentioned in Section 2.4. The dip-dist test could not reject unimodality at *p* = 0.05. The folding test accepted unimodality at *p* = 0.0001 (Φ = 29.06). Further 10, 000 the random Gaussian projections of the progression vectors onto 1-D were all unimodal at *p* = 0.05 with Bonferroni and Benjamini-Hochberg corrections. As we did for Baseline Scores, we also tested progression vectors for unimodality using SigClust. The results, available in Supplementary Materials, support the hypothesis of unimodality.

### 2.13 Neighborhoods for Progression Vectors

Using the same methodology as used for baseline scores, we hierarchically clustered the 33-D progression vectors. Similar to baseline vectors, Ward’s metric, silhouette scores, and gap statistics (see Figure 5b-d) did not reveal any natural clusters. Using these metrics, we selected six neighborhoods named *P*_1_ − *P*_6_ as sufficiently high in number to examine different regions of the data but still a reasonable choice according to the previously mentioned statistics (Figure 5b shows the dendrogram, details of the choice of six clusters are in the Section 4.9). The multidimensional scaling projections of *P*_1_ − *P*_6_ are shown in Figure 5c. The figure also shows the population in each neighborhood, as well as bar graphs for the mean values of the B+R, Tremor, and Axial progression subscores.

Table 4 shows the mean and std. dev. of TMS scores, B+R, Tremor, and Axial subscores in *P*_1_, …, *P*_6_. We carried out paired t-tests comparing the TMS, B+R, Tremor, and Axial subscore progression vectors in neighborhoods *P*_1_ − *P*_6_. The lines below the mean (std. dev.) numbers show in bold which neighborhood had significantly different means (at *p* = 0.05 with Benjamini-Hochberg correction for multiple comparisons). Table 4 shows that each neighborhood differs from every other neighborhood on some subscore. For example, *P*_2_ differs from *P*_1_, *P*_3_, *P*_4_, *P*_6_ in the B+R subscore, and from *P*_4_ in the Tremor subscore. This suggests that progression vectors are significantly heterogeneous.

**Table 4:** Comparison of the 6 neighborhoods identified from the projection vectors. For numeric variables: means and standard deviations of participants in each neighborhood. Bold-faced neighborhoods below a mean indicate that the above mean is significantly different from the mean of the bold-faced neighborhood.

| Neighborhood | $P_1$ | $P_2$ | $P_3$ | $P_4$ | $P_5$ | $P_6$ |
| --- | --- | --- | --- | --- | --- | --- |
| $\Delta H\&Y$ | 0.15(0.2)<br>P1 P2 P3 P4 P5 P6 | 0.29(0.25)<br>P1 P2 P3 P4 P5 P6 | 0.37(0.24)<br>P1 P2 P3 P4 P5 P6 | 0.3(0.26)<br>P1 P2 P3 P4 P5 P6 | 0.44(0.23)<br>P1 P2 P3 P4 P5 P6 | 0.59(0.3)<br>P1 P2 P3 P4 P5 P6 |
| MDS-UPDRS Part 1 |  |  |  |  |  |  |
| $\Delta Total$ | 2.98(4.06)<br>P1 P2 P3 P4 P5 P6 | 2.75(4.84)<br>P1 P2 P3 P4 P5 P6 | 4.43(4.88)<br>P1 P2 P3 P4 P5 P6 | 3.24(4.25)<br>P1 P2 P3 P4 P5 P6 | 3.96(5.0)<br>P1 P2 P3 P4 P5 P6 | 4.71(6.21)<br>P1 P2 P3 P4 P5 P6 |
| MDS-UPDRS Part 2 |  |  |  |  |  |  |
| $\Delta Total$ | 0.69(2.32)<br>P1 P2 P3 P4 P5 P6 | 1.95(2.91)<br>P1 P2 P3 P4 P5 P6 | 2.78(3.82)<br>P1 P2 P3 P4 P5 P6 | 1.5(2.51)<br>P1 P2 P3 P4 P5 P6 | 3.7(3.02)<br>P1 P2 P3 P4 P5 P6 | 4.9(4.37)<br>P1 P2 P3 P4 P5 P6 |
| MDS-UPDRS Part 3 |  |  |  |  |  |  |
| $\Delta TMS$ | 2.65(1.42)<br>P1 P2 P3 P4 P5 P6 | 6.07(1.77)<br>P1 P2 P3 P4 P5 P6 | 10.07(1.63)<br>P1 P2 P3 P4 P5 P6 | 7.44(2.16)<br>P1 P2 P3 P4 P5 P6 | 13.06(2.18)<br>P1 P2 P3 P4 P5 P6 | 11.59(3.96)<br>P1 P2 P3 P4 P5 P6 |
| $\Delta B+R$ | 2.15(1.32)<br>P1 P2 P3 P4 P5 P6 | 4.72(1.32)<br>P1 P2 P3 P4 P5 P6 | 8.06(1.19)<br>P1 P2 P3 P4 P5 P6 | 4.8(1.55)<br>P1 P2 P3 P4 P5 P6 | 8.84(2.03)<br>P1 P2 P3 P4 P5 P6 | 7.83(3.29)<br>P1 P2 P3 P4 P5 P6 |
| $\Delta Tremor$ | -0.16(0.94)<br>P1 P2 P3 P4 P5 P6 | 0.42(0.66)<br>P1 P2 P3 P4 P5 P6 | 0.41(0.84)<br>P1 P2 P3 P4 P5 P6 | 1.78(1.04)<br>P1 P2 P3 P4 P5 P6 | 2.97(1.12)<br>P1 P2 P3 P4 P5 P6 | 0.28(0.7)<br>P1 P2 P3 P4 P5 P6 |
| $\Delta Axial$ | 0.53(0.46)<br>P1 P2 P3 P4 P5 P6 | 0.75(0.47)<br>P1 P2 P3 P4 P5 P6 | 1.35(0.5)<br>P1 P2 P3 P4 P5 P6 | 0.68(0.38)<br>P1 P2 P3 P4 P5 P6 | 1.0(0.49)<br>P1 P2 P3 P4 P5 P6 | 3.24(0.79)<br>P1 P2 P3 P4 P5 P6 |
| Progression Vector Length | 1.28 | 1.58 | 2.39 | 1.87 | 2.90 | 3.05 |

### 2.14 Heterogeneity of progression vector length and direction

The analysis in the previous section is based on movement subscores. A different perspective is obtained if we take each 33-D progression vector and analyze its length and direction in the 33-D space. We compared the length of the progression vectors in each neighborhood. The mean length of the progression vectors in each neighborhood is shown in the last row of Table 4. A paired t-test of the lengths of progression vectors in *P*_1_, …, *P*_6_ reveals that the mean lengths are all different at *p* = 0.05 (with Benjamini-Hochberg correction). *P*_6_ has the largest mean length, while *P*_1_ has the smallest. Further, *P*_3_ and *P*_4_ have larger progression lengths than *P*_2_ and *P*_5_ respectively.

Dividing the progression vector by its length projects it onto the unit sphere (see Figure 1a5 for an illustration of this idea). This unit length vector represents the direction of the progression vector. The means of the unit vectors in each neighborhood are visualized in 2-D in Figure 6 using multidimensional scaling. The visualization in Figure 6 suggests that *P*_2_, *P*_3_, and separately *P*_4_, *P*_5_, are similar in direction. To test this statistically, we used the integrated likelihood ratio test (ILRT) for mean directions [42]. The ILRT for mean directions was selected because it is known to perform well in high dimensions [42]. The ILRT for mean direction compares two groups of unit vectors for equality of mean. Using the ILRT to compare equality of mean in every pair of neighborhoods shows that the mean unit vectors in *P*_2_, *P*_3_ have a similar directions (*p* = 4.6*e* − 11). The same is true for *P*_4_, *P*_5_ (*p* = 3.4*e* − 11).

**Figure 6:**
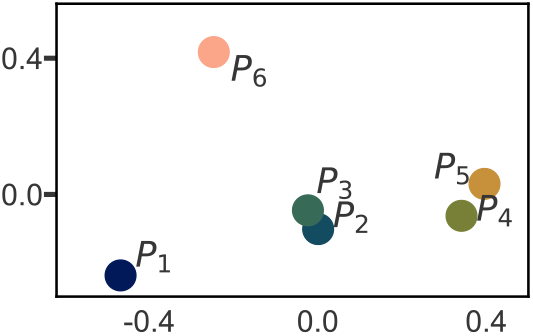
Mean directions of each neighborhood visualized in 2-D (2-dimension) using multidimensional scaling.

### 2.15 Additional Characteristics of *P*_1_, *…, P*_6_

We compared the age and sex of the participants in *P*_1_, …, *P*_6_ (Table 5). Participants in *P*_6_ are nearly a decade older (mean age of 70.2) than participants in the other groups (mean age of 60.6 for *P* 1 − *P* 5). There are no statistically significant differences in age among *P*_1_ − *P*_5_. *P*_2_ − *P*_5_ have more men than women. *P*_1_, however, has more women than men (31 men/34 women) but comparing *P*_1_ to the rest of the population *P* 2 − *P* 6 using a two-sample z-test for proportions does not yield a statistically significant difference after correcting for multiple comparisons (*p* = 0.009 *> p* = 0.0083).

**Table 5:**
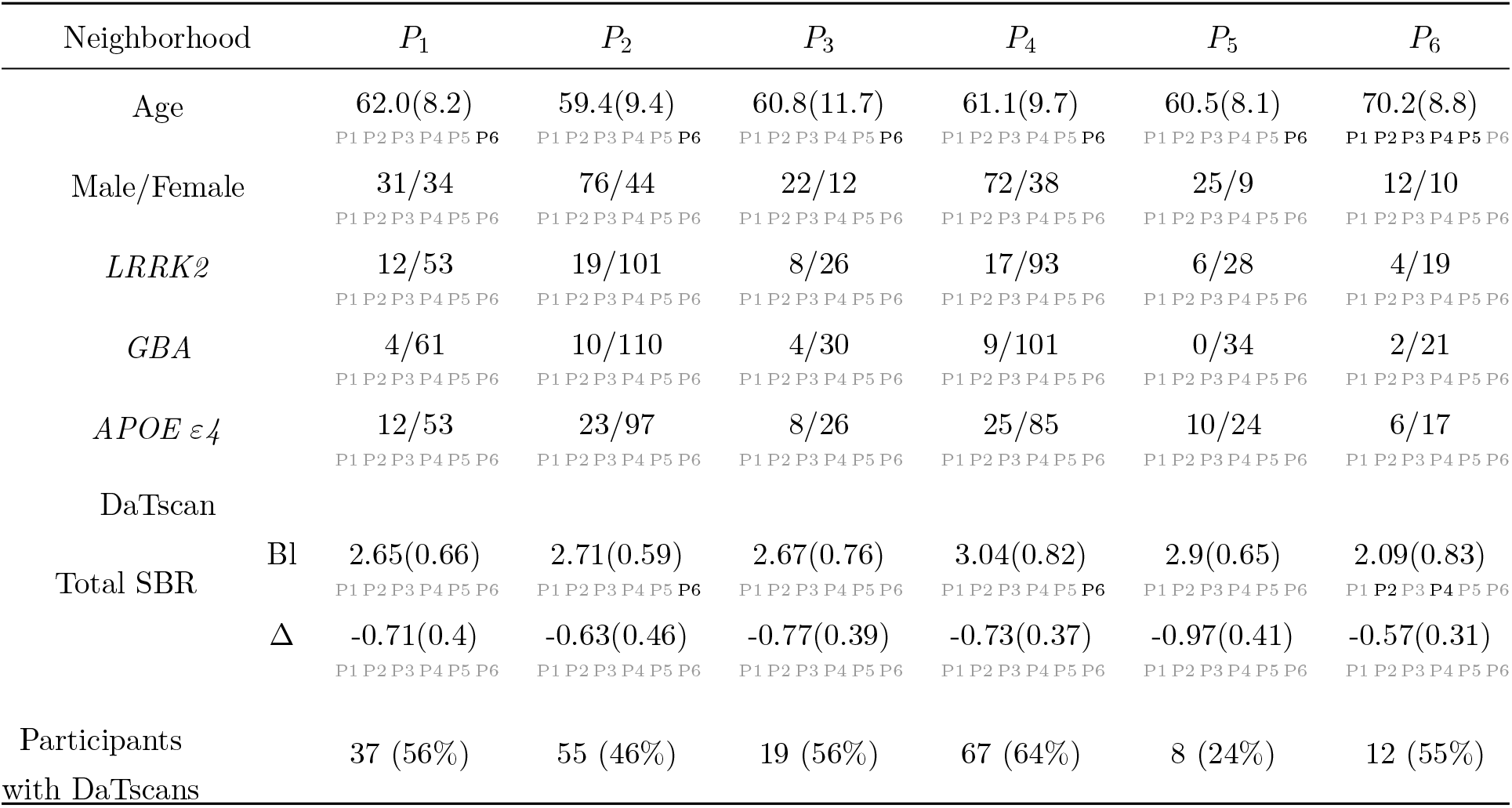
Comparison of age, sex, genetics, and DaTscan imaging in *P*_1_ − *P*_6_ identified from the projection vectors. For numeric variables: means and standard deviations of participants in each neighborhood. For genetics: numbers indicate YES/NO counts. For *LRRK2* and *GBA*, this is the presence of a pathogentic mutation. For *APOE ε4*, this is the presence of *APOE ε4*. Bold-faced neighborhoods below a value indicate that the above value is significantly different from the value of the bold-faced neighborhood.

We next compared the prevalence of pathogenic genetic variants for *LRRK2* and *GBA* and labels of *APOE ε4* alleles for each participant (Table 5). Rates of pathogenic genetic mutations were not found to be significantly different between any progression neighborhoods.

Finally, we compared DaTscan images of participants in each neighborhood. DaTscan images were compared using the total striatal binding ratio (SBR) as described in Methods Section 4.12. The SBR was calculated at baseline and at year four for 198 of the 386 participants who had DaTscan images from both times. The fraction of subjects in each neighborhood for whom DaTscans were available is shown in the last row of Table 5. Except of *P*_5_ the neighborhoods have more or less similar fraction of subjects with DaTscans.

To account for variation in the actual time between the baseline imaging and the 4-year imaging, the 4-year change in SBR was linearly interpolated. At baseline, we found that the total SBR in *P*_6_ was significantly lower than in many other neighborhoods (the mean total SBR in *P*_6_ was 2.09 while the mean total SBR for *P*_1_ − *P*_5_ ranged from 2.65 − 3.04. See Table 5).

### 2.16 Summary of progression vector heterogeneity in movement scores

In summary, the following picture emerges of progression vector heterogeneity (see Figure 7a for a multidimensional scaling visualization of the progression vectors in the neighborhoods.):

1. Neighborhoods *P*_2_, *P*_4_ contain over 60% of the participants. When additionally combined with *P*_3_, *P*_5_, these neighborhoods contain 88% of the population. Thus these four neighborhoods contain majority of the population. *P*_2_, *P*_3_, *P*_4_, *P*_5_ have the following characteristics:
  - *P*_2_ and *P*_3_ have similar mean progression directions, but the mean progression vector in *P*_3_ is bigger than the mean progression vector in *P*_2_. That is, *P*_3_ is a faster progressing version of *P*_2_. A similar argument holds for *P*_4_ and *P*_5_; *P*_5_ is a faster version of *P*_4_.
  - *P*_2_ has moderate progression of B+R subscores while tremor and axial subscores progress relatively slowly. *P*_4_ also exhibits moderate progression of B+R subscores, but in addition, it exhibits moderate progression of the tremor subscore.
  - All of the above can be summarized by saying that progression for most of the patients can be explained as variability along two “axes” (see Figure 7a). The magnitude of tremor progression varies along one axis, while progression vector length (i.e the speed of progression) varies along the other.
2. *P*_6_ and *P*_1_ are the exceptional neighborhoods. Each of these neighborhoods contains less than 10% of the population.
3. 1. *P*_6_ has the longest progression vectors (fastest progression speed) of all participants. Progression vectors in *P*_6_ show large changes in B+R and axial subscores. However, tremor subscores are virtually unchanged. *P*_6_ also has the oldest population, and significantly lower SBRs.
4. *P*_1_ has the smallest progression vectors (slowest speed). *P*_1_ has mildly changing B+R subscores, while other subscores do not change significantly.
5. There are no genetic mutation differences between the neighborhoods.

**Figure 7:**
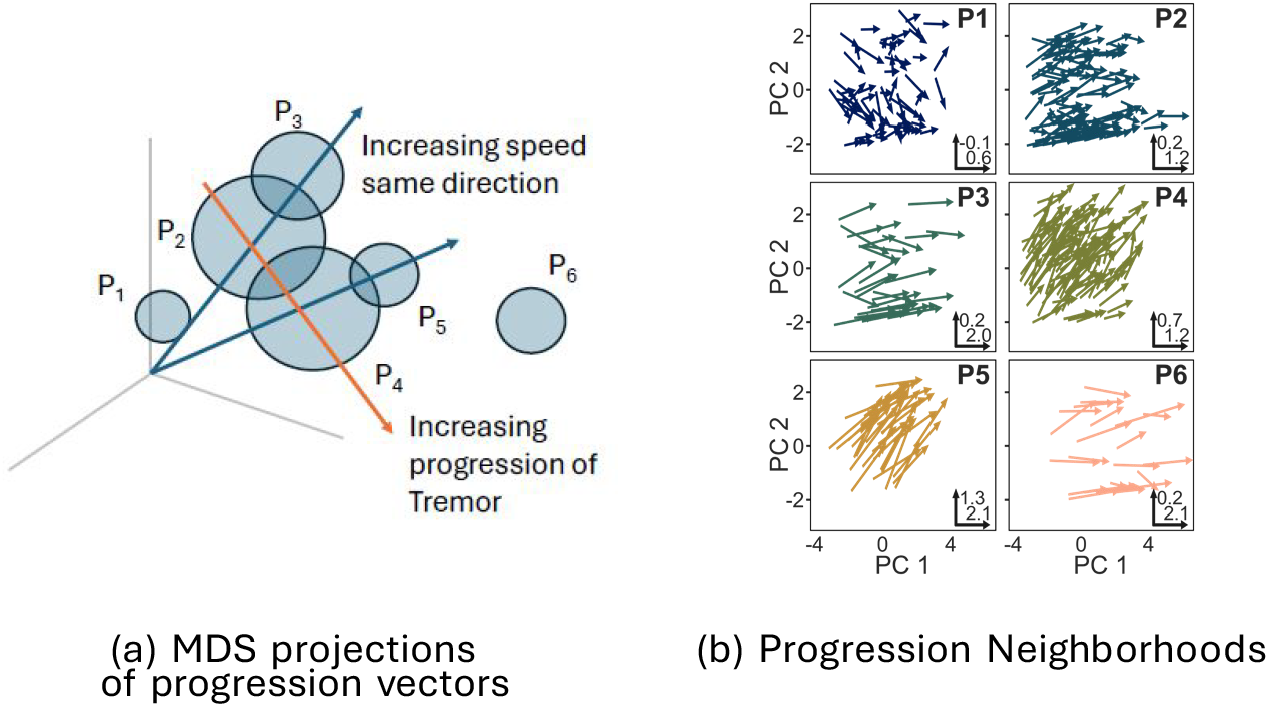
Visualization of progression vectors and their neighborhoods.(a) Multi-dimensional scaling (MDS) projections of progression vectors in the neighborhoods. (b) An illustration of the relative locations of progression vector neighborhoods. The means of *P*_2_, *P*_3_ and *P*_4_, *P*_5_ lie along lines through the origin. PC (principle component)

## 3 Discussion

Parkinson’s disease presents and progresses heterogeneously, yet there is no clear consensus on a taxonomy for this heterogeneity. Possible reasons for this are that studies may be underpowered for the amount of noise in the data, or that the ordinal nature of PD clinical scores is not taken into account. In some cases, the non-Markovian nature of progression may hinder the analysis. We attempted to overcome these limitations by using data from a large study, by explicitly denoising the longitudinal motor scores of the participants, and analyzing baseline scores and progression vectors. The analysis of progression vectors could be simplified to a linear analysis because PD progression is slow in the early stage and can be approximated linearly as discussed in the Results section. Extending our analysis to longer time durations may require a more sophisticated approach.

Visualizing baseline and progression scores shows that both are heterogeneous, but there do not appear to be any obvious and distinct clusters in either. As mentioned earlier, there is no generally acceptable statistical test of unimodality, but the dip-dist, the folding test, and the random projections strongly suggest that baseline scores and progression vectors may be better thought of as continuously variable rather than as subtypes. This conceptualization is consistent with a recent call for analyzing PD heterogeneity without explicit subtypes [8].

The analysis of denoised baseline and progression vectors shows that the latter are not linearly predictable from the former (PLS and PCR are linear models). That is, motor-score progression is not linearly Markovian from baseline. In principle, supplementing baseline motor scores with other information (e.g. demographics or DaTscans) may make progression predictable. It is also possible that a nonlinear model might demonstrate increased predictability. A third and intriguing possibility is that while the entire 33-D progression vector is not predictable, some meaningful low-dimensional function of it might be. This possibility is suggested by the study [43] where gait, freezing of gait, and instability scores are summed into an ambulatory capacity measure, which is then shown to be predictable from baseline.

Finally, different baseline score neighborhoods have a very similar distribution of age, sex, and genetic mutations with the exception of *LRRK2. LRRK2* mutations have a higher rate in *B*_2_, *B*_4_ compared to *B*_1_, *B*_3_. *B*_2_, *B*_4_ also appear to have lower tremor scores than *B*_1_, *B*_3_. One possible explanation for this is the slower progression of PD for *LRRK2* mutation as noted in [44].

In contrast to baseline scores, progression vectors offer more meaningful information: most of the PD participants are contained in neighborhoods *P*_2_ − *P*_5_. The progression in these neighborhoods can be explained by using two axes. The progression in Tremor subscores varies along one axis, while the overall speed of progression varies along the other.

Two smaller subsets of the population are contained in *P*_1_ and *P*_6_, The relatively high baseline scores and very gradual progression of *P*_1_ do not match any previously reported subtype of PD progression. *P*_1_ does have the highest percentage of women (although not statistically significant), a factor that might be related to the slower progression. But note that previous comparisons of differences in PD progression in men and women have not yielded conclusive results [45, 46].

Interestingly, despite slow progression in *P*_1_, the DaTscans SBRs for this group showed no significant differences when compared to *P*_2_ − *P*_5_. Participants such as these – with very slow symptom progression but with decreasing total SBR typical of PD – may partially explain why some studies [47–50] found inconsistent and low correlations between motor symptoms and DaTscans.

*P*_6_ is perhaps the most distinct of the six neighborhoods, with the oldest population and the fastest progression of axial symptoms and H&Y stage. *P*_6_ bears similarity to the late-onset subtype proposed previously [14, 15, 51]. *P*_6_ is consistent with the idea that late onset is more severe [26]. *P*_6_ is also comparable to the PIGD-D clinical subtype [2, 3, 13], which is characterized by higher initial axial symptoms and more rapid progression. However, we do not find baseline axial subscores to be higher in *P*_6_ compared to *P*_1_ − *P*_5_. Additionally, higher initial H&Y scores in *P*_6_ suggest that patients in *P*_6_ may also be further along in their disease trajectory at the time of enrollment in the study.

While P1 and P6 are intriguing subpopulations, they do contain small number of subjects. It is possible that with a larger study, the characteristics of P1 and P6 might change. In this context, we also want to point out a possible selection bias in *P*_6_. As we noted above, *P*_6_ contains the oldest population, and given our selection criterion (at least 5 visits in 4 years) it is possible that some subjects who might otherwise fall into category are excluded (members of the older population may not have 5 visits in 4 years).

Regarding the relation of genetic mutations and PD progression, previous literature has shown a weak protective relationship between *LRRK2* mutations and speed of motor symptom progression [21, 23, 44]. But there are conflicting results [52] which suggest that *LRRK2* mutations may be associated with more severe PIGD symptoms. Previous research comparing *GBA* mutations and motor symptoms also have mixed results, some [23] finding no relationship between the two while others [22] show that GBA was associated with more rapid motor-symptom progression. Finally, some have found *APOE ε4* to have no effect on motor symptom progression [20], while others have found a correlation with faster progression but only after 5+ years [19].

For all three of these genetic risk factors (*LRRK2, GBA*, and *APOE ε4*), we found no significant differences between any of the progression-vector neighborhoods. In part, this may be due to the relatively small number of participants with pathogenic versions of each mutation (29(7.5%) for *GBA*, 66(17.1%) for *LRRK2*, 84(21.8%) for *APOE ε4*) and that we only used data from 4 years of early stage progression. Note that Tan et al. [53] also found no significant relationship between any genetic mutations and motor-symptom progression.

One limitation of our study is that it focuses on relatively early-stage PD. It is possible that later stages of the disease may be more Markovian and less heterogeneous. Moreover, all of our analysis is retrospective, and some effort is needed to make it predictive. Another limitation of the study is that over 96% the participants in the study are Caucasian and all are US-based. There is evidence that ethnicity and nationality have an effect on PD progression. For example, at first diagnosis, the African American population in the US appears to present with a later stage of the disease [54–56]. It is possible that the longitudinal progression and the heterogeneity of the disease may also differ in this subpopulation. Other mono-ethnicity studies show differences in the prevalence of various PD subtypes around the world [57]. Turning to nationality, there is evidence to suggest that phentotyping of PD in closely related ethnicities differ based on geographic location [58]. Currently, there is no data set comparable to the size and complexity of PPMI to address these issues. However, we note that our methodology would be applicable without change to such datasets.

In conclusion, our data-driven denoising and clustering approach provides novel insights into the heterogeneity of baseline and progression characteristics of PD and sheds light on the root causes of inconsistencies in previous reports. Our methodology can serve as an analytical linchpin tying together the biological and clinical characteristics of the disease anchored in an in-depth quantitative understanding of its inherent heterogeneity. Our findings also have implications for recent efforts to develop biological staging frameworks of PD for research purposes [59, 60] as well as for outcome assessments in clinical trials using disease-modifying therapies. For example, our results suggest that denoising is important in assessing PD progress (staging) and the effect of therapy on it. Our results also suggest that any clustering of baseline or of progression measures should be accompanied by clear statistical evidence that the clustering is statistically valid.

## 4 Methods

### 4.1 Inclusion and Exclusion of Participants

Data were obtained by the Parkinson’s Progression Markers Initiative (PPMI, http://www.ppmiinfo.org) [38]. The criteria for inclusion and exclusion in the PPMI PD cohort are available in [38]. MDS-UPDRS Part III assessments were conducted by PPMI at the following monthly intervals: 0, 3, 6, 9, 12, 18, 24, 30, 36, 42, 48, 54, 60 and then annually. We excluded MDS-UPDRS assessments that were done in the “ON medication” state as defined by PPMI (*<* 12 hours since last dose of LDOPA or LDOPA equivalent). Assessments which were conducted remotely, or which only had partial data, or which occurred after a participant had received deep brain stimulation surgery were also excluded. From the remaining, we selected participants who satisfied all of the following criteria: a clinical assessment within the first six months of their participation, a clinical assessment at least 4 years or longer after baseline, and at least 4 clinical assessments in total. This resulted in a pool of 386 participants, which we used in our analysis.

### 4.2 Genetics

Genetic testing for *LRRK2, GBA*, and *APOE ε4* was performed as part of the PPMI study. Labels of pathogenic mutations for both *LRRK2* and *GBA* provided for each participant by PPMI were used. Participants with either one or two *APOE ε4* alleles as determined by PPMI were considered positive for *APOE ε4* in our analyses.

### 4.3 Denoising of MDS-UPDRS Part III Scores

The denoising method is specifically designed for use with MDS-UPDRS Part III questions [37]. The method is implemented in MATLAB and is available for download (https://github.com/tagarelab/Ordinal-Time-Series-Denoising). As explained below, the method is a hierarchical, maximum-likelihood procedure.

The main idea behind the method is illustrated in Figure 8. Suppose there are *N* subjects, with each subject represented by an ordinal time series of measurements. An ordinal regression model (regression with time) can be fit to each time series. The ordinal regression models the probability distribution of ordinal scores and their evolution in time. The regression model is explained in more detail below. We assume that the mean of the distribution represents the underlying “signal”, and the variability around the mean represents “noise”. The temporal change in the mean thus indicates any progression of the score. Note that the mean of an ordinal distribution is continuous-valued and not discrete, and thus the progression of the mean can be viewed as a smooth “interpolation” of the noisy discrete ordinal scores.

**Figure 8:**
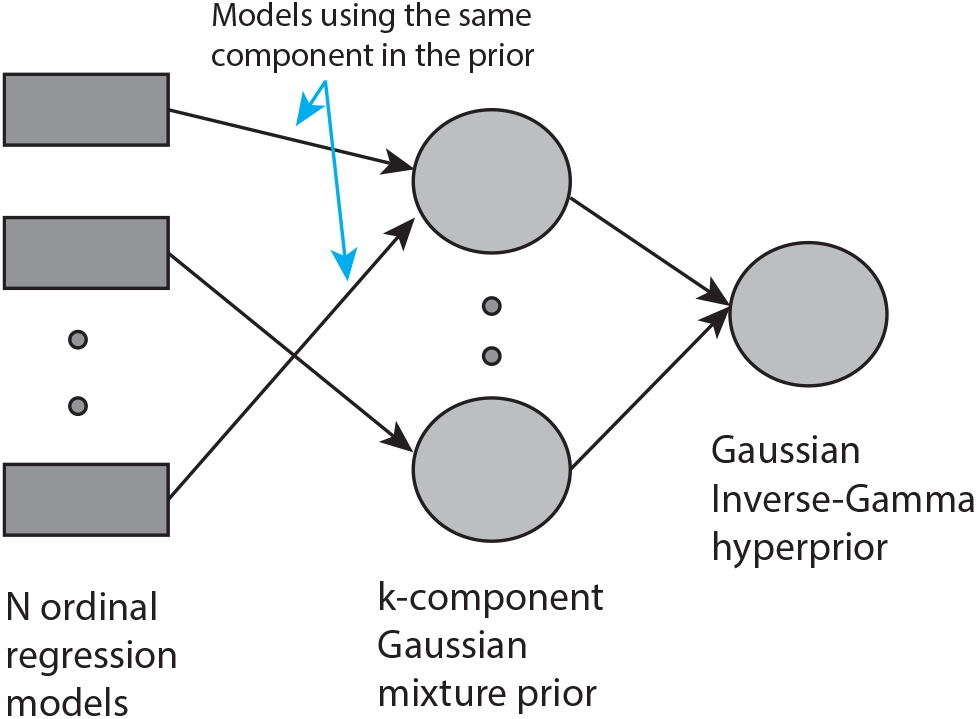
Ordinal Time Series Denoising Model. Given *N* time series the model uses *N* ordinal regression models, one for each time series. The parameters of the regression model come from a *k*-component Gaussian mixture prior. Each component (mode) of the prior represents similarly evolving time series. The parameters of the components of the prior share a conjugate hyperprior, representing the overall progression of all time series.

When the ordinal time series is short, the estimate of the probability distribution may not be reliable. However, if there are sufficiently large number of subjects, it is likely that several subjects have similar (but not identical) time series. These subjects can “pool” their data by assuming that the coefficients of their ordinal regression come from a single prior. More generally, assuming a mixture prior with *k* components and associating each subject’s regression with one component enables the regressions to draw statistical power from each other (see Figure 8). All time series whose models share a component in the prior are similar. They are also distinct from time series which share any other component. By changing the variance of the Gaussian components, the model adapts to tightly clustered time series heterogeneity (using small component variance) or more continuously varying time series heterogeneity (using large component variance). The prior components themselves have parameters and these parameters are assumed to come from a single hyperprior which represents the overall progress of the disease in the entire population.

To be specific, the ordinal regression model we use is the a variant of the proportional-odds model [61]. Briefly, the model is as follows (this description is adapted from [37]):

Suppose *y* is an ordinal random variable taking values in ordinal categories { 0,…, *C* − 1}. A model for the probability distribution of *y* is

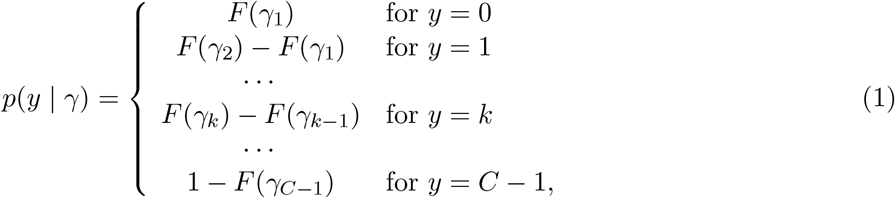

where *F* is the cumulative distribution function (cdf) of the standard normal density and *γ* = (*γ*_1_, *…, γ*_*C* − 1_) *∈ R*^*C* − 1^ is a parameter of the distribution subject to the constraint *γ*_1_ ≤ *γ*_2_ ≤ *γ*_3_ ≤ *…* ≤ *γ*_*C* − 1_. It is useful to implicitly impose these constraints by a change of variables. Define a map Γ which maps *u* = (*u*_1_, *…, u*_*C* − 1_) to *γ* = (*γ*_1_, *…, γ*_*C* − 1_) as follows:

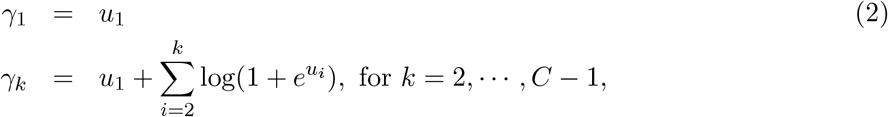

Then the ordinal probability model of equation (1) can be written as *p*(*y* |Γ(*u*)) with no constraints on *u*.

The above model extends to a time-series model as follows: suppose *Y* = (*y*_1_, …, *y*_*T*_) is a discrete ordinal time series with *y*_*i*_ ∈ {0,, *C* − 1} for observed at time *t*_*i*_ ∈ [0, 1] for *i* = 1,…, *T*, with *t*_*i*+1_ *> t*_*i*_. Note that no assumption is made about the time series being uniformly sampled in time (i.e. *t*_*i*_ need not be equal to (*i* − 1)*/*(*T* − 1)). Then,

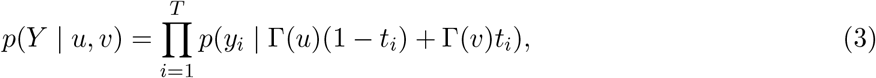

where *u, v ∈* ℝ^*C* − 1^ generate the thresholds *γ* that divide the probabilities at the start and end of the time interval [0, 1]. This model effectively assumes a linear change over time of the *u* value used in *p*(*y* | Γ(*u*)). The model can be made more complex by making the change non-linear, but for slowly progressing diseases such as PD, a linear change is sufficient. To proceed, let *θ* = (*u, v*).

Suppose there are *N* subjects, with time series 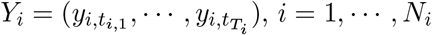. Assuming that the *i*th subject has the subject’s own parameter *θ*_*i*_ = (*u*_*i*_, *v*_*i*_),

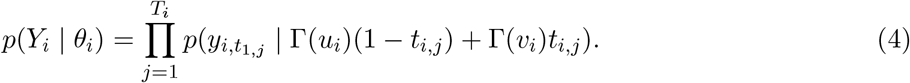

Note that this formulation allows each subject to have a time series with different number of points (number of points in the time series = *T*_*i*_), and the data in the series need not be uniformly sampled in time.

As mentioned above, we set the prior for *θ*_*i*_ to a normal mixture with *K* components:

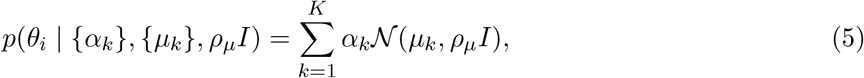

where *µ*_*k*_ are the means of the components, and *ρ*_*m*_*u* is the variance (assumed diagonal and equal for all components). The *µ*_*k*_’s and *ρ*_*µ*_ are hyperparameters, and we use the following hyperprior for them:

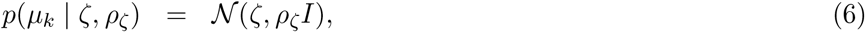

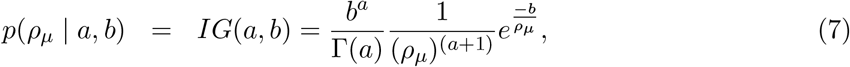

where *IG* is the inverse-gamma distribution. The hyperprior parameters *a, b, ρ*_*ζ*_ are fixed, by a crossvalidation procedure described in detail in [37]. The cross-validation procedure also establishes, *K*, the number of clusters in the prior of equation (5).

Grouping together all parameters to be estimated as *ϕ* = (*{θ*_*i*_}, {*µ*_*k*_}, *ρ*_*µ*_, {*α*_*k*_}, *ζ*), we get:

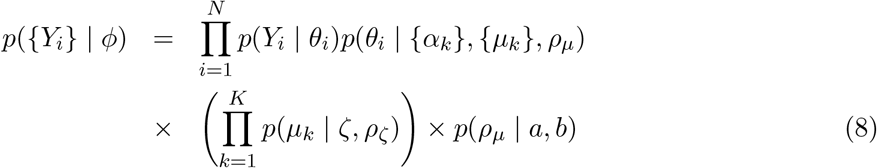

from which the maximum-likelihood estimates of *ϕ* are obtained as 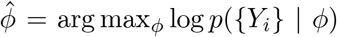. The maximum-likelihood estimates are easily calculated by the EM-algorithm whose details are given in [37].

The maximum likelihood estimates of 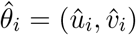 give the estimated mean for the *i*th subject at time *t* as:

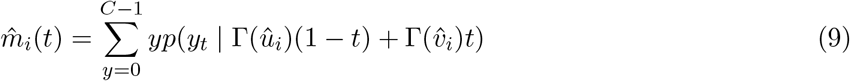

The estimated mean is the denoised signal.

Mathematical details of the model are available in [37]. That publication also evaluates the method with simulations and real MDS-UPDRS Part III scores. Simulations with data representing MDS-UPDRS Part III scores show that the method is superior to alternative filtering methods. Further experiments with real MDS-UPDRS Part III data shows that the method does not overfit: the distribution of the log-likelihood of training and test data is very similar.

### 4.4 MDS-UPDRS Part III Score Processing

For each participant, the more severely affected side was determined by comparing sums of the denoised scores for all left-side and all right-side scores at baseline. The more severely affected side at baseline was considered more severely affected for all future visits.

To conveniently summarize results in this paper, we grouped and summed MDS-UPDRS Part III questions into three categories according to the cardinal signs the questions address as follows:

- Bradykinesia and Rigidity (B+R): 3.2, 3.3, 3.4, 3.5, 3.6, 3.7, 3.8 (18 scores)
- Tremor: 3.14, 3.15, 3.16, 3.17, 3.18 (10 scores)
- Axial: 3.9, 3.10, 3.11, 3.12 (4 scores)

MDS-UPDRS question 3.1, which addresses speech did not fit any of these categories, was not included in any of the sums, but was included in the vector representing baseline and progression scores.

### 4.5 Dimensionality Reduction and Statistical Analyses

Multidimensional scaling (MDS) was performed in Python using the Scipy and Scikit-learn libraries. Multidimensional scaling (MDS) was then used to reduce the 33-D progression vectors to 2-D for visualization.

### 4.6 Kernel Density Estimation

Kernel density estimation was used to visualize the density of points in 2-D space. We used the *gaussian*_*kde* function from the Scipy Python package.

### 4.7 Tests for unimodality

Hartigan’s dip-test is the classic unimodality test [39]. We used the “diptest” package in R for this test.

The dip-dist test [40] calculates Euclidean distances from each point to all other points. Dip-dist tests the unimodality of the distances at each point by using Hartigan’s dip-test. Thus Hartigan’s test is applied multiple times, once at every point. We correct for multiple testing by using the more conservative Bonferroni correction and the less conservative Benjamini-Yekutieli correction. The latter is used because distances from a point to all other points is likely to be correlated.

The folding test [41] folds data with respect to a center. The variance of the folded data is smaller than the variance of the original data. When the center is appropriately chosen (see [41] for details) the reduction in the variance is larger for multimodal data compared to unimodal data. The folding test is based on this idea. The test is available as an R package (Rfolding). Our tests were carried out using this package.

If the data is clustered and the clusters are linearly separable, then an appropriate projection of the data onto a 1-D subspace can be multimodal. We projected data onto 10, 000 random subspaces and applied Hartigan’s dip test to each projection, while correcting for multiple comparisons using Benjamini-Hochberg and Bonferroni corrections.

### 4.8 PLS and PCR

Partial Least Squares Regression (PLS) and Principal Component Regression are classic methods in statistics. We used implementations of both methods in the Python package Scikit-learn.

### 4.9 Hierarchical Clustering

Baseline and progression vectors were clustered using Scikit-learn’s Agglomerative Clustering [62]. Agglomerative clustering begins by treating each data point as a cluster and then sequentially merging clusters so as to reduce the within-cluster variances. The process ends when all data have been merged into a single cluster. The merging process is represented as a tree (the dendrogram), with the individual data points as leaf nodes, and a single cluster as the root node. At any level in the tree, the effectiveness of the clustering is measured by Ward’s metric, which is the sum of the within-cluster variances of the data. Figure 9a shows this quantity as a function of number of clusters for baseline scores. Figure 9b shows the same for progression vectors. Since the baseline and progression vector data are shown to be unimodal, there are no natural clusters. There is no single, generally accepted method for selecting the optimal number of clusters so we examined several different metrics. The first was Ward’s metric which is shown in the dendrogram. We also calculated the silhouette coefficients (Figure 4c and Figure 5c) and gap statistics (Figure 4d and Figure 5d) for different numbers of clusters [63]. Both were calculated using the agglomerative clustering results and the Python package, Scikit-learn. For the silhouette score, it is common to pick the number of clusters at the “elbow” of the plot – the number of clusters before the values plateau. Although neither graph has a clear “elbow”, 4 clusters and 6 clusters occur before the silhouette scores plateau. For the gap statistic, it is common to select the number of clusters for which there is a large gap between the gap statistic for *n* and *n* − 1 clusters.

**Figure 9:**
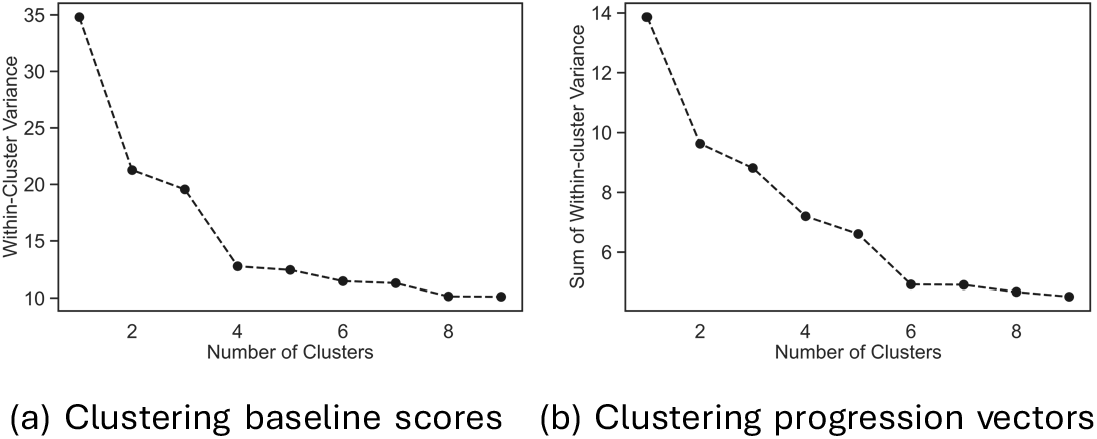
Plots of the total within-cluster variances for each number of clusters using agglomerative clustering for (a) baseline scores and (b) progression vectors. These values correspond to the heights of divisions in the dendrogram.

Neighborhoods were identified with Scikit-learn’s Agglomerative Clustering [62]. Ward’s method was used to form linkages. We selected the number of clusters by examining the dendrogram and selecting a cut-point with large distances between clusters. More specifically, this method stops clustering at a point where further merging of clusters would substantially increase the total within-cluster variance (the metric used for Ward’s method). To make meaningful statistical comparisons, we only considered sets of clusters where all clusters had populations above 10. This resulted in 4 clusters for baseline vectors and 6 for the progression vectors.

### 4.10 Pairwise comparisons between neighborhoods

Means of numerical quantities between neighborhoods were assessed with the Student’s t-test with a significance threshold of *p* = 0.05. Binary labels such as sex and genetic variants were compared between neighborhoods with a two sample Z-test for equality of proportions. All pairs of neighborhoods were compared. To account for multiple comparisons, the Benjamini-Hochberg correction was used with a 5% false discovery rate.

### 4.11 Integrated likelihood ratio test (ILRT)

We used the variant of the integrated likelihood ratio test (ILRT) for comparing mean directions of groups of vectors described in [42]. This test was chosen for its performance in high-dimensional cases (*d >>* 3) and its ability to deal with groups with low counts. Direction was defined as the 33-D unit vector with the same direction as the progression vector for all 33 MDS-UPDRS Part III scores.

### 4.12 DaTscan Analysis

DaTscan images are available for PPMI participants at baseline and then annually for up to the next 5 years, with possible missing scans. Our analysis included participants who had DaTscans at baseline and at year 4. This criterion was satisfied by 199 of the 386 participants. DaTscans are already aligned by PPMI to the Montreal Neurological Institute’s (MNI) Atlas. We obtained regions for the bilateral caudates and putameni from the MNI Atlas. The DaTscans were normalized using the occipital region and then mean striatal binding ratios (SBRs) were calculated for the two caudates and two putameni. These four values were added together to get the total SBR. Since not all scans were exactly 4 years apart, SBR values were linearly interpolated.

## Data Availability

The data used in this article is publicly accessible and available from PPMI (www.ppmi-info.org).

## Code Availability

The denoising software used in this article is available from GitHub (https://github.com/tagarelab/Ordinal-Time-Series-Denoising).

## Acknowledgments

This work was supported by the NIH (NINDS) under the Grant R01NS107328. The data used in the preparation of this article were obtained from the Parkinson’s Progression Markers Initiative (PPMI) database (www.ppmi-info.org/data). For up-to-date information on the study, visit (www.ppmi-info.org). PPMI – a public-private partnership – is funded by the Michael J. Fox Foundation for Parkinson’s Research and funding partners. The full list of funding partners can be found at (www.ppmi-info.org/about-ppmi/who-we-are/study-sponsors/).

## Author Contribution Statement

JDK, HDT, and ST jointly developed the data analysis methodology. JDK analyzed the PPMI data. ST helped with the interpretation of the results. All authors contributed to the writing. All authors have read and approved the final manuscript.

## Competing Interests

The authors declare no competing interests.

## Notes

### Competing Interest Statement

The authors have declared no competing interest.

