## Supplement for "Denoised MDS-UPDRS Part-III Scores Yield New Patterns of Progression Heterogeneity in Early Stage Parkinson’s Disease"

### 1 Principal Component Analysis

We disregarded the longitudinal nature of the data, took all scores of all participants as raw data, and carried out principal component analysis (PCA). Assessing whether the significant principal components stay similar or not after denoising indicates whether denoising eliminated noise or signal. Separate PCAs were performed for the raw and denoised scores and the top principal components (PCs) are then compared.

The loadings and explained variance of the top four PCs for raw and denoised data (shown in Supplementary Figure 1) are similar. This suggests that denoising does not have a significant effect on the PCs. That is, denoising does not significantly affect the signal.

The first PC has positive loadings for all non-tremor questions (B+R and axial). The second PC has positive loadings for tremor questions. Together, these two PCs explained over 50% of total variance. These results are similar to previous reports which performed factor analysis on the MDS-UPDRS Part III. The top two factors identified in [1] were gait, posture, and speech (36.8% explained variance) and tremor (15% explained variance). Similarly, [2] also performed factor analysis and proposed a two-factor model: non-tremor and tremor.

### 2 Cluster Analysis

To further test whether the baseline MDS-UPDRS Part III scores and the corresponding progression vectors were better characterized as a single cluster or two clusters the methodology in [3] was applied to both the baseline and progression vector data using the SigClust package in R. For both the baseline MDS-UPDRS Part III scores and the progression vectors, SigClust did not provide significant evidence against the null hypothesis of a single cluster. Specifically, the baseline analysis yielded a non-significant p-value ($p = 0.165$), and the progression-vector analysis likewise yielded a non-significant p-value ($p = 0.680$), indicating that a two-cluster solution ($k = 2$) was not favored over a one-cluster solution ($k = 1$) in both cases.

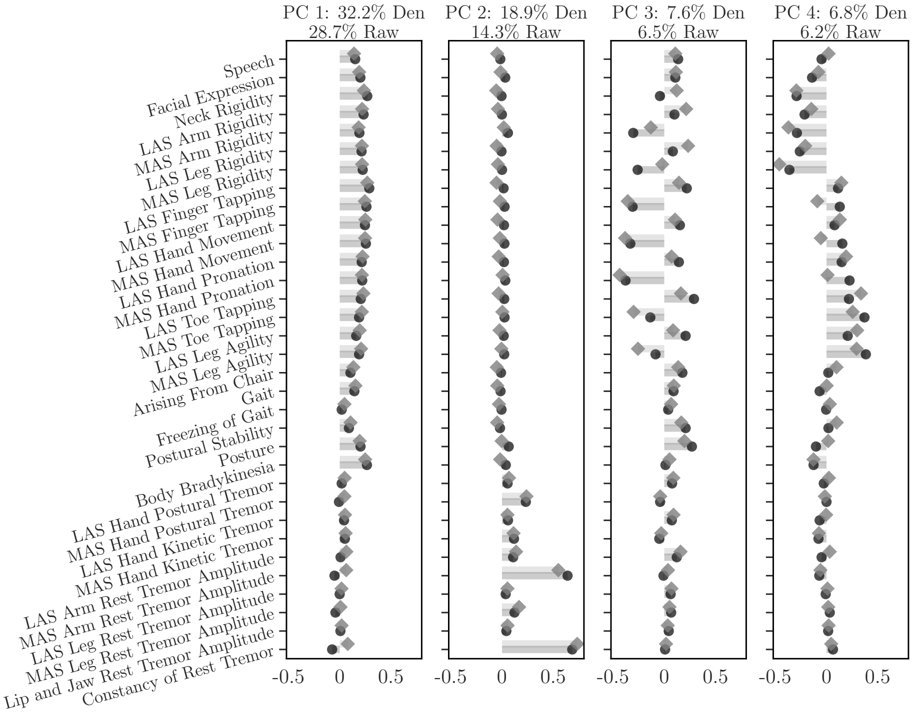


**Supplementary Figure 1.** Comparison of principal component loadings before (gray diamonds) and after denoising (black squares) for the first four principal components (PCs). Explained variance percentages for each PC are listed above the respective graphs.

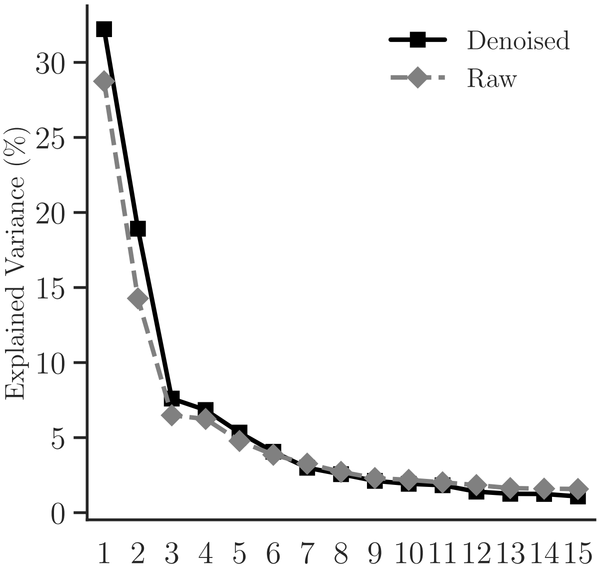


**Supplementary Figure 2.** Comparison of the explained variance due to the top 15 principal components before (gray diamonds) and after denoising (black squares).

[2] Michelle Hyczy de Siqueira Tosin, Christopher G Goetz, Sheng Luo, Dongrak Choi, and Glenn T

Stebbins. Item response theory analysis of the mds-updrs motor examination: tremor vs. nontremor items. *Movement Disorders*, 35(9):1587–1595, 2020.

[3] Y. Liu, D. N. Hayes, A. Nobel, and J. S. Marron. Statistical significance of clustering for high-dimension, low-sample size data. *Journal of the American Statistical Association*, 103(483): 1281–1293, Sep. 2008.
